# Viral burdens are associated with age and viral variant in a population-representative study of SARS-CoV-2 that accounts for time-since-infection related sampling bias

**DOI:** 10.1101/2022.12.02.518847

**Authors:** Helen R. Fryer, Tanya Golubchik, Matthew Hall, Christophe Fraser, Robert Hinch, Luca Ferretti, Laura Thomson, Anel Nurtay, Lorenzo Pellis, George MackIntyre-Cockett, Amy Trebes, David Buck, Paolo Piazza, Angela Green, Lorne J Lonie, Darren Smith, Matthew Bashton, Matthew Crown, Andrew Nelson, Clare M. McCann, Adnan Mohammed Tariq, Rui Nunes Dos Santos, Zack Richards, The COVID-19 Genomics UK (COG-UK) consortium, David Bonsall, Katrina A. Lythgoe

## Abstract

In this study, we evaluated the impact of viral variant, in addition to other variables, on within-host viral burdens, by analysing cycle threshold (Ct) values derived from nose and throat swabs, collected as part of the UK COVID-19 Infection Survey. Because viral burden distributions determined from community survey data can be biased due to the impact of variant epidemiology on the time-since-infection of samples, we developed a method to explicitly adjust observed Ct value distributions to account for the expected bias. Analysing the adjusted Ct values using partial least squares regression, we found that among unvaccinated individuals with no known prior infection, the average Ct value was 0.94 lower among Alpha variant infections, compared those with the predecessor strain, B.1.177. However, among vaccinated individuals, it was 0.34 lower among Delta variant infections, compared to those with the Alpha variant. In addition, the average Ct value decreased by 0.20 for every 10 year age increment of the infected individual. In summary, within-host viral burdens are associated with age, in addition to the interplay of vaccination status and viral variant.

## Introduction

The SARS-CoV-2 epidemic in the United Kingdom (UK) has been characterised by the appearance of a series of distinct viral variants that, in order of emergence, include the B.1.177 lineage, and the Alpha (B.1.1.7 lineage), Delta (B.1.617.2 lineage) and Omicron (BA.1, BA.2, BA.4 and BA.5 lineages) variants. Explaining their successive abilities to spread, the Alpha, Delta and Omicron variants have been estimated to have a transmission advantage of 43-100% [1-3], 60-70% [4] and 52% [5] compared to their preceding variant. The underlying causes of these differences are unclear, but could include differences in within-host viral burdens [6], infectious periods, or the per-virion probability of between-host transmission. In turn, these could be influenced by many factors [7], including changes in virus attachment to human cells and the continuous interplay of population acquisition of immunity and the emergence of immune escape variants [8, 9]. In this study, we compare within-host viral burdens of different viral variants by analysing nose and throat swabs collected as part of the UK’s nationally representative SARS-CoV-2 surveillance study [10, 11].

A number of studies have compared viral burdens between the Alpha variant and predecessor variants (Supplementary Table 1)[12-18] with mixed findings. For example, two detailed longitudinal surveys of a small number of infected individuals have suggested that viral burdens are similar among the variants [16, 17]. However, a much larger, but less intensive study of viral burdens at symptom onset has identified higher viral burdens among individuals infected with the Alpha variant, compared to those with a predecessor lineage [15]. The impact of later variants on viral burdens has also been studied [11, 15, 16, 19], indicating higher viral burdens associated with the Delta variant compared to the Alpha variant, among vaccinated individuals [11] in one survey, but no difference in viral burdens among these variants in another [16]. The study design and cohorts used to investigate viral burdens have varied and this may explain the different findings. In addition to the differences in sample sizes and sampling frequency, the study populations have varied. Some have been based upon testing symptomatic individuals or their close contacts [12, 14, 15] and have thereby excluded some asymptomatically infected individuals, who make up an estimated 40% [20] of infections. Others have focussed on a specific group of people, with examples being hospitalized individuals [12] and persons associated with a professional sporting league [16]. Methods to identify variants have also varied, with some surveys using Spike gene target failure (SGTF) [12, 14, 15] during PCR testing or sample date [11] to classify the viral variants, whereas other have used whole genome sequencing [13, 16, 17].

The Office for National Statistics (ONS) COVID-19 Infection Survey (CIS) is a large household-based surveillance study based in the United Kingdom [10, 11]. We analysed data from the CIS to investigate the impact of viral variant on viral burdens. The survey randomly selects private households on a continuous basis from address lists and previous surveys to provide a representative UK sample. Individuals were asked to provide information that included demographics, symptoms, and vaccination details. As part of the survey, nose and throat swabs were collected and tested for SARS-CoV-2 using RT-PCR, and, if positive, individuals with a cycle threshold (Ct) less than 30 were sequenced using whole genome sequencing. Since the Ct value of a sample is inversely correlated with log^10^(viral burden) of that sample [21], this study design enables viral burdens to be investigated. Although the accuracy with which sampled viral burdens from nose and throat swabs informs viral burdens occurring throughout the body is unclear [22], this study does allow for investigation into viral burdens in a manner that avoids biases associated with samples from symptomatic individuals or small studies of particular demographic groups.

The survey is simultaneously a cross sectional survey of the population through time and a longitudinal survey of individuals, with individuals sampled approximately weekly during the first month following enrolment and then monthly thereafter. This weekly or monthly sampling leads to uncertainty in the time-since-infection of positive samples. In addition, the different epidemiological trajectories of the variants mean that the distribution of time-since-infection for each variant at any given time can be skewed depending on when the samples were collected. For example, if a variant is increasing in prevalence, a cross sectional sample will contain more individuals with that variant who are earlier on in their infection compared those who are later on in their infection[23]. Because within-host viral burden trajectories are asymmetric, with the peak in viral load closer to the start of infection than to the end [16], this can affect the sampled distribution of viral burdens and complicate comparisons between viral variants. The impact of SARS-CoV-2 epidemiology on sampled Ct values is sufficiently strong for its shifts to be inferred from changes in Ct values measured over time [23, 24].

We are unaware of any published studies comparing viral burdens associated with viral variants from a large population-representative surveillance survey that directly estimates the impact of variant-specific epidemiological trajectories. Here, we address this gap by developing a methodology that directly estimates the combined impact of variant-specific within-host viral burden and epidemiological trajectories on randomly sampled viral burdens. We apply this methodology to data from the CIS to investigate the impact of a range of factors, including variant, vaccination status, and age, on viral burdens, as measured by Ct values. As many countries move towards implementing SARS-CoV-2 surveillance surveys, the concepts and methodologies described here will be valuable for informing public health decisions.

## Results

We analysed RT-qPCR SARS-CoV-2 positive samples from the CIS that were sequenced at Oxford (sampled between 27/09/20 and 17/06/21) or Northumbria (sampled between 20/09/21 and 19/01/22) and had a Ct≤30. These samples cover the period of the epidemic that includes part of the B.1.177 wave, the full Alpha wave, part of the Delta wave, and part of the BA.1 Omicron wave. The lineages of sampled sequences were identified from sequence data (see methods for details) and only the samples that could be classified as either B.1.177, or the Alpha, Delta or BA.1 (Omicron) variants of concern (VoCs) were analysed. Of a total 10586 and 24232 sequences obtained from samples sent to Oxford and Northumbria in which a lineage could be assigned, 3256 (31%) and 477(2%) respectively were not from these lineages and were excluded from further analysis.

### A framework to infer epidemiologically adjusted Ct values

To enable us to compare viral burdens between different viral variants, we developed a framework that adjusts observed Ct values to account for the different epidemiological trajectories of different viral variants (see methods). In brief, variant-specific incidence rates for each of the major variants in the sample data (B.1.177, Alpha, Delta and BA.1 Omicron) (Figure 1a) were inferred by combining estimates of total SARS-CoV-2 incidence rates in England (www.ons.gov.uk/peoplepopulationandcommunity/healthandsocialcare/conditionsanddiseases/datasets/coronaviruscovid19infectionsurveydata) with estimates of the proportion of incident infections with each variant, as inferred from the COVID-19 infection consortium data repository (www.cogconsortium.uk/). These data were used rather than the equivalent estimates available directly from CIS to prevent the introduction of a time lag between incidence and prevalence into our study. The variant-specific incidence rates were combined with normally distributed infection periods to estimate how the expected distribution of time since infection from randomly sampled individuals changes over calendar time for each of the variants. For each PCR positive sequenced sample in our analysis, the expected distribution of the time since infection corresponding to its variant and sample date was identified and truncated to account for expected bounds, where these could be determined by previous positive or negative samples from the same individual.

**Figure 1.**
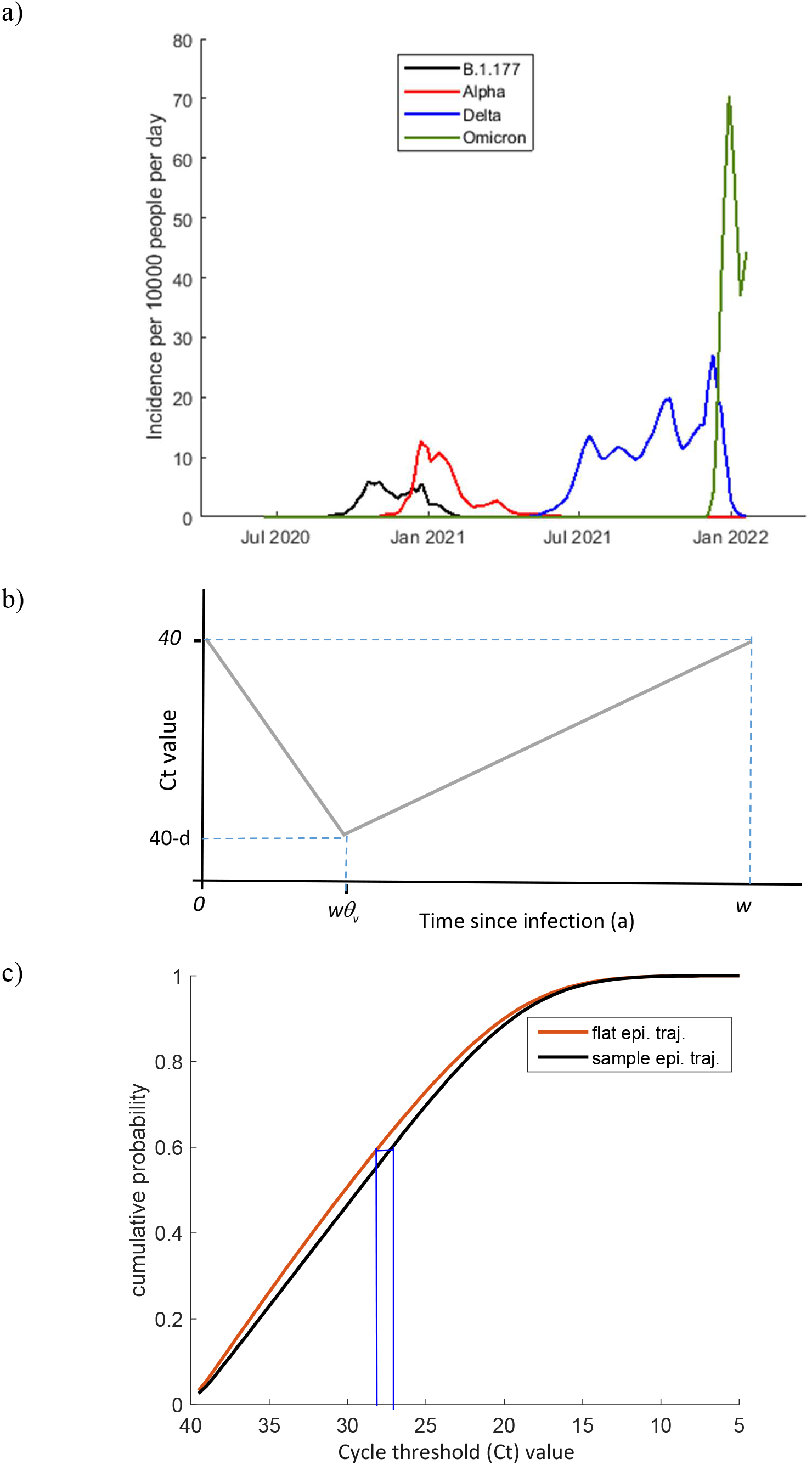
A method for estimating epidemiologically adjusted Ct values. a) Inferred daily incidence with the B.1.177 lineage and the Alpha, Delta and BA.1 Omicron variants between July 2020 and January 2022 in the UK. These were estimated to equal the product of the total daily incidence and the fraction of incident infections of that variant. b) Within-host Ct trajectories were assumed to be valley shaped, with infected period (width) *w*, and depth *d*. The valley trough was estimated to be a fraction *θ*_*v*_ across the width. c) Adjusted Ct values were inferred by first estimating the cumulative probability distribution of Ct values based upon the sample date and the known epidemiological trajectory of the sample variant and identifying the percentile at which the observed Ct value falls within this distribution. Second, the cumulative probability distribution of Ct values under an assumption of a flat epidemiological trajectory was estimated and the Ct value at the selected percentile was identified.

For each sample, we next estimated an expected distribution of Ct values. This was achieved by assuming that within-host Ct values are described by a piecewise, valley-shaped trajectory (Figure 1b) with depth (viral burden peak) and width (infected period) taken from normal distributions. The timing of the valley trough (peak viral burden) was fixed at a chosen fraction across the width. The parameters describing these metrics were estimated from an alternative data source [16]. However, the mean maximum valley depth (mean peak viral burden) was iteratively inferred, and other parameters – including the timing of peak viral burden – were varied during sensitivity analyses. For each sample, an adjusted Ct value was then inferred by finding the percentile of the observed Ct value among the expected Ct distribution and selecting the Ct at the corresponding percentile in an expected Ct distribution, calculated from a flat epidemic trajectory (Figure 1c).

### Ct values from early and late during the Alpha wave are more closely aligned after epidemiological adjustment

Since we had data spanning the full epidemiological trajectory of the Alpha wave in the UK, we determined the impact of our method when applied to data collected at different stages during its trajectory. We applied the adjustment to Alpha-variant samples collected from individuals who were unvaccinated and had not been identified as being spike-antibody positive prior to infection (n=3413). By splitting the samples according to sample date into two equally sized sets (early-phase and late-phase) we visualised how the timing of sampling during the epidemiological trajectory impacted observed Ct values (Figure 2).

**Figure 2.**
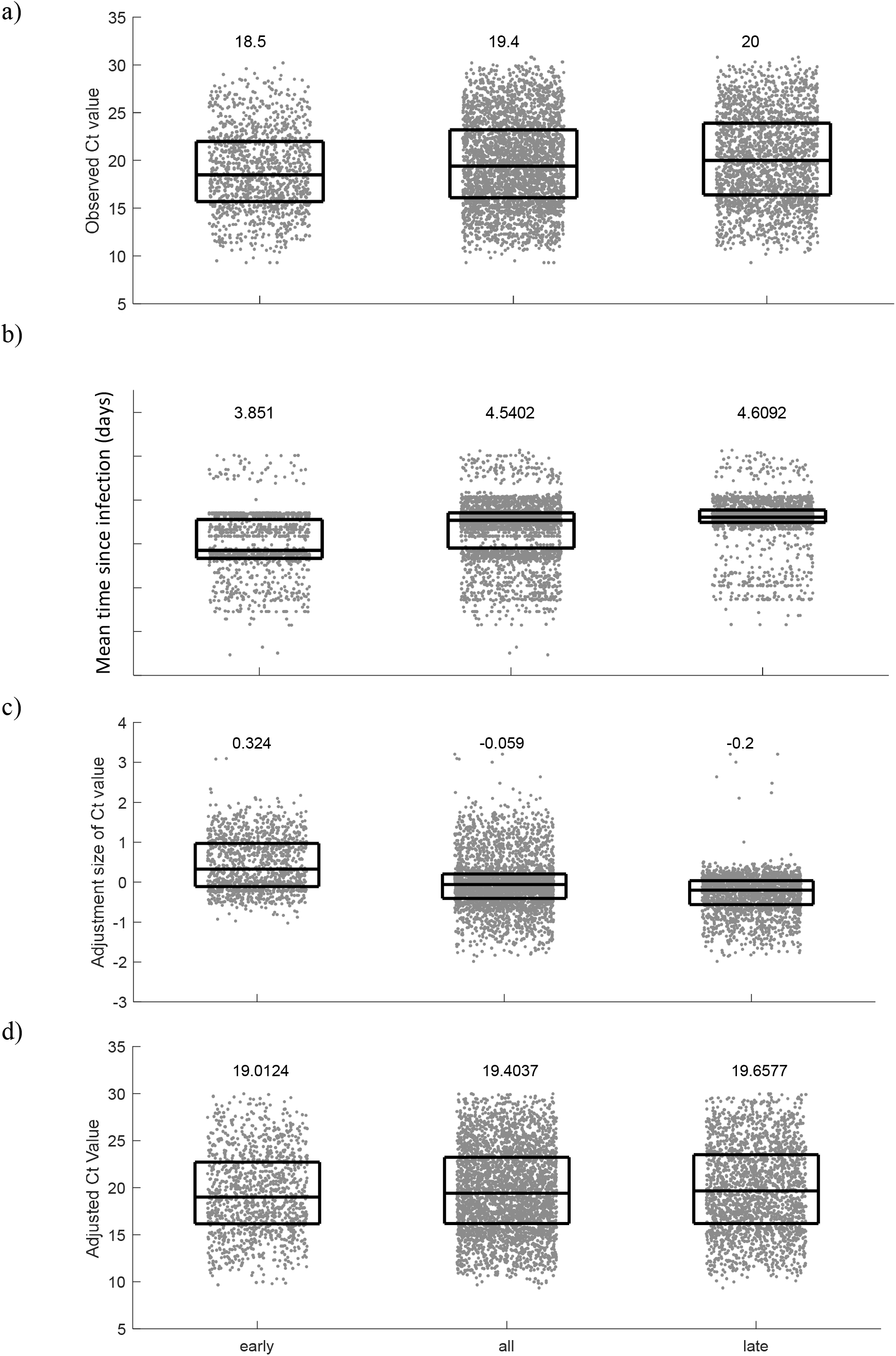
Epidemiological adjustment results in more closely aligned estimates of mean viral burden from samples taken early and late during the Alpha wave. Samples that correspond to Alpha-variant infections in individuals who are unvaccinated and have not been identified as being antibody positive prior to infection are split according to sample date. Four metrics are applied to data from the early phase, all phases and the late phase. In each panel, median and interquartile ranges are overlaid onto individual data points. a) The observed Ct values are, on average, higher for late phase, compared to early phase samples. b) The estimated mean time since infection is, on average, longer for late-phase, compared to early-phase samples. c) The Ct adjustment size is, on average, positive for early phase samples, negative for late phase samples and negligible when all data are considered. d) On average, the adjusted Ct values relating to the early and late phase are more closely aligned than the observed Ct values. However, adjusted values remain, on average, higher in late-phase, compared to early-phase samples.

The median of the unadjusted Ct values was lower for early-phase samples than for late-phase samples (Figure 2a), consistent with the expected impact of the epidemiological effect. For each sample, our method estimates a probability distribution for the time since infection for that sample, based upon the sample variant, sample collection date, and, where available, the dates of recent positive and negative sample within the same infection. The mean time since infection derived from each of these distributions is plotted in Figure 2b. On average the mean time since infection is longer among the late-phase compared to early-phase samples. Because Ct values are, on average, lower in early infection compared to late infection (Figure 1b), the adjustment acted in the opposite direction and increased the Ct values of early-phase samples, but decreased the Ct values of late-phase samples (Figure 2c).

When the epidemiological adjustment was applied to the Ct values, the adjusted distribution of Ct values for the early-phase and late-phase were more closely aligned compared to the unadjusted values (Figure 2d). For comparison, the application of the method to data from the whole Alpha wave is also shown (middle column in Figure 2), revealing that the net adjustment applied to the full set of samples is negligible. This emphasises the value of using the epidemiological adjustment when samples are only available for part of the epidemiological trajectory of a variant, such as during the emergence phase of a new variant.

### The asymmetry of the within-host viral trajectory impacts comparisons

Our framework highlights that the combined impact of the shape of the within-host viral trajectory and the epidemiological stage of a variant can affect viral burdens measured at the population level. Plausible changes to our assumption of the mean infected period have only a small impact upon the adjusted values (Figure 3a), whereas plausible changes to the fractional position of the viral burden peak across this period have a much bigger effect on the adjusted values (Figure 3b) (although absolute changes are still relatively modest compared with variability between individuals). The closer the peak viral burden is to the start of infection, the greater the epidemiological correction applied to samples selected from just early on or just late on during the Alpha wave. This can be understood by noting that in a random sample, early-phase samples have, on-average, shorter times since infection than late-phase samples and the greater the asymmetry of the within-host viral burden, the greater the difference in expected viral burdens between infections in the earlier or later phases of infection (Figure 3c).

**Figure 3.**
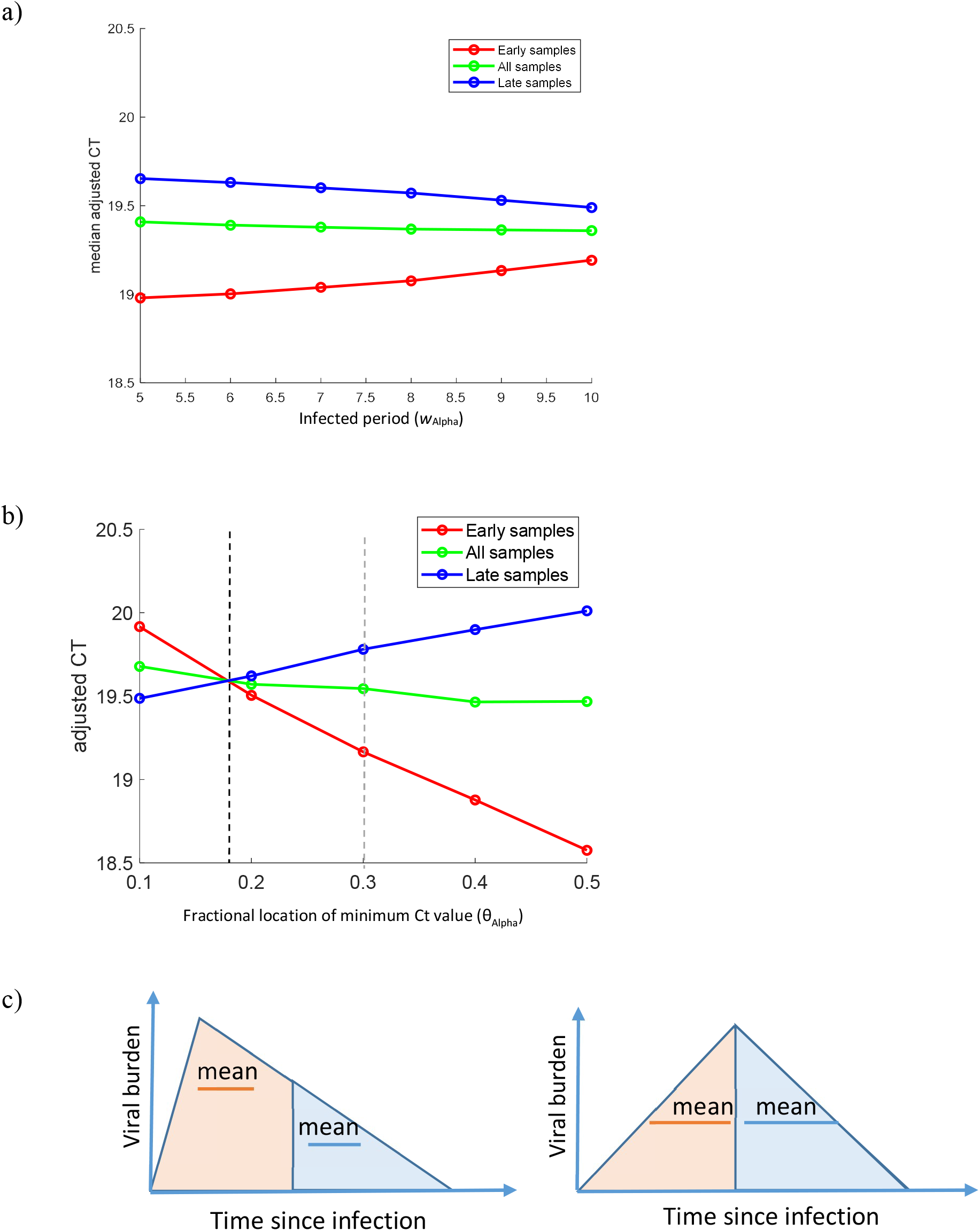
The epidemiological stage and the asymmetry of the within-host viral trajectory impact the Ct adjustment size. In panels a) and b) samples that correspond to Alpha-variant infections in individuals who are unvaccinated and have not been identified as being antibody positive prior to infection are split according to sample date. The medians of the adjusted Ct values are plotted for early samples (red), late samples (blue) and all samples (green) under different assumptions about the asymmetry and the mean width of the within-host viral burden trajectory. In panel a) the infected period is varied under an assumption that the viral burden trajectory is skewed towards the start of infection (*θ*_Alpha_=0.3). This shows that Ct values are lower (viral burdens are higher) amongst samples taken earlier on during infection, but vary to only a limited degree with changes in the mean infected period (*w*_Alpha_). In panel b) the fractional location of the peak viral burden, *θ*_Alpha_, is varied under the assumption that the mean infected period is 8 days (*w*_Alpha_=8). This shows that the asymmetry of the within-host viral burden trajectory measurably impacts the adjusted Ct values and that the early and late-phase Alpha variant samples are most closely aligned when *θ*_Alpha_=0.18. Panel c) highlights how when the within-host trajectory is skewed towards earlier during infection, viral burdens sampled during early infection will on average be higher than those sampled later on in infection.

In calculating the adjusted Ct values for samples with the Alpha variant (Figures 2d) we assumed that peak viral burden occurs at a fraction 0.3 across the infected period, based upon prior data from 103 individuals [16]. It is noteworthy that using this parameter estimate the median adjusted Ct value remains slightly higher for late-phase, compared to early-phase samples. This can be visualised by comparing the red and blues lines shown in Figure 3b at a value *θ*_Alpha_=0.3 along the x-axis (grey dashed vertical line). By identifying the intersection of these two lines, it is possible to show that the median adjusted Ct values of the early-phase and late-phase samples are equal when the asymmetry of the within-host trajectory is increased, such that *θ*_Alpha_=0.18 (black dashed vertical line). Arguably, changing this parameter estimate so that the peak is closer to the start of infection than we have assumed may provide a better estimate of its true value compared to the one that we derived from a prior study. However, there are other explanations for higher viral burdens (lower Ct values) in the early-phase samples. Because the CIS has not intensively sampled participants throughout the pandemic, but rather conducted a large round of recruitment in September-October 2020, meaning that many participants at the start of the Alpha wave were still undergoing more regular – approximately weekly – follow-up, they may genuinely have been sampled closer to the start of infection in the early phase than the later phase. Second, CIS tested antibodies in only ∼15% participants prior to the Alpha wave, so we cannot rule out that some samples come from individuals who had had a prior infection and that the number of such individuals has increased over the duration of the Alpha wave. It is thus credible that more intensive sampling and lower population levels of immunity present earlier on in the Alpha wave could contribute to the pattern of lower adjusted Ct values in early-phase compared to late-phase samples.

### Investigating factors associated with viral burdens

We investigated whether factors, including viral variant, are associated with adjusted Ct values sampled in the CIS and sequenced at Oxford or Northumbria Universities using partial least squares regression (PLS). Samples sequenced at Oxford were collected between 27^th^ September 2020 and 17^th^ July 2021, and cover the period of the epidemic that includes part of the B.1.177 wave, the full Alpha and part of the Delta wave. Samples sequenced in Northumbria were collected between 20^th^ September 2021 and 19^th^ January 2022 and cover part of the Delta wave and part of the BA.1 Omicron wave. We have analysed the samples sequenced from the two centres separately so that differences in sequencing protocols and inclusion criteria do not affect our results.

Adjusted Ct values for samples from these two centres are shown in Figure 4, categorised according to sample date (Figure 4a and 4d), participant age (Figure 4b and 4e), and a combination of prior exposure category and variant (Figure 4c and 4f). Using partial least squares regression analysis, we assess the impact of sample date, sex, first vaccine dose product (AstraZeneca, Pfizer) and prior exposure category (no known prior exposure, prior exposure without vaccination, 1 vaccine dose, 2 vaccine doses, 3 vaccine doses) on adjusted Ct values. In addition, within each prior exposure we assess the impact of variant (B.1.177, Alpha, Delta and BA.1 Omicron).

**Figure 4.**
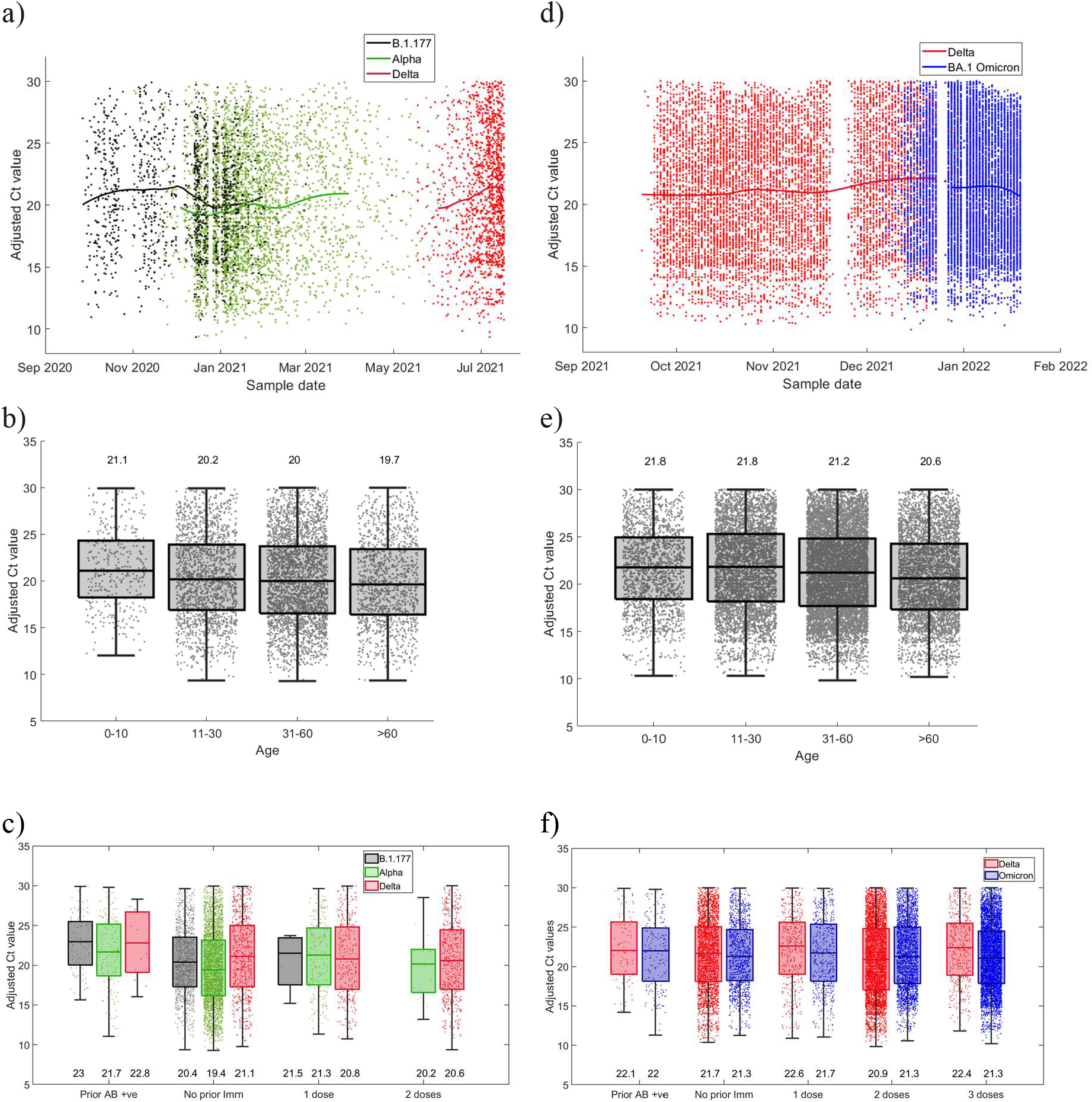
Adjusted Ct values plotted against different factors. For samples sequenced at Oxford (a, b and c) and at Northumbria (d, e and f), adjusted Ct values are plotted against different factors. Panel a) and d) show a LOESS fit (smoothing parameter=0.55) of adjusted Ct values over sample date, categorised by variant. Panels b) and e) show box and whisker plots of adjusted Ct values by age category. Panels c) and f) show box and whisker plots of adjusted Ct values by prior vaccination and/or infection, by variant. Horizontal lines represent the median and interquartile range. Parameter values used in these calculations are listed in table 3.

Prior to application of the PLS regression model, we investigated multicollinearity among predictor variables, by calculating variance inflation factor (VIF) values (supplementary table 2). A VIF> 3 can be considered an indicator of moderate multicollinearity and a VIF>5 an indicator of strong multicollinearity. Among the Oxford samples, we found that sample date was highly multicollinear with other predictors (VIF 7.5). This is intuitively clear from the strong temporal separation of samples with the Alpha and Delta variants, as observed in Figure 4a. Because we wanted to assess the potential impact of variant, and since it is not possible to independently assess the effect of sample date and variant, we removed sample date from the regression model of the Oxford samples. In the interpretation of the subsequent analysis, it is therefore important to recognise that factors that have not explicitly been included in the regression, but correlate with calendar time, cannot be ruled out as being predictive of viral burdens. For the Northumbria samples, although the sample date VIF value of 3.4 indicates only a moderate degree of collinearity, for consistency we similarly removed sample date from the regression.

**Table 1.** Beta scores and variance in projection (VIP) values for the partial least squares analysis of samples sequenced in Oxford and Northumbria.

| Samples | Oxford |  | Northumbria |  |
| --- | --- | --- | --- | --- |
| Result | Beta score | VIP | Beta score | VIP |
| *Included in Model* |  |  |  |  |
| **Age (in years)** | -0.013 | 1.38 | -0.026 | 2.15 |
| **Prior immunity**<br>*Ref = AB -ve and unvaccinated* |  |  |  |  |
| AB +ve | 3.29 | 0.42 | 0.33 | 0.24 |
| Vaccinated | 1.42 | 1.33 | 0.47 | 0.88 |
| **Variant (in AB -ve and unvaccinated)**<br>*Ref = Alpha (Oxford), Delta (Northumbria)* |  |  |  |  |
| B.1.177 | 0.94 | 1.42 |  |  |
| Delta | 1.06 | 1.52 |  |  |
| BA.1 Omicron |  |  | -0.14 | 0.60 |
| **Variant (in AB+ve)**<br>*Ref = Alpha (Oxford), Delta (Northumbria)* |  |  |  |  |
| B.1.177 | 0.78 | 0.35 |  |  |
| Delta | -2.91 | 0.22 |  |  |
| BA.1 Omicron |  |  | -0.07 | 0.23 |
| **Variant (in vaccinated)**<br>*Ref = Alpha (Oxford), Delta (Northumbria)* |  |  |  |  |
| B.1.177 | -0.31 | 0.05 |  |  |
| Delta | -0.34 | 1.02 |  |  |
| BA.1 Omicron |  |  | -0.15 | 0.88 |
| **Vaccine dose in vaccinated**<br>*Ref = 1 dose* |  |  |  |  |
| 2 doses | 0.01 | 0.75 | 0.05 | 0.82 |
| 3 doses |  |  | 0.51 | 0.81 |
| **Vaccine product**<br>*Ref = Pfizer* |  |  |  |  |
| AstraZeneca | -0.09 | 1.02 | -0.09 | 1.00 |
| *Not included in model* |  |  |  |  |
| **Sex**<br>*Ref = Female* |  |  |  |  |
| Male | -0.13 | 0.35 | -0.13 | 0.63 |

**Table 2.** Data used in the study.

| Variable | Description |
| --- | --- |
| $t_{ijk}$ | Sample date of the $k$th sample of the $j$th infection of the $i$th individual |
| $\tilde{t}_{ij}$ | Sample date of the last negative before the first positive of the $j$th infection of the $i$th individual |
| $c_{ijk}$ | Observed Ct value of the $k$th sample of the $j$th infection of the $i$th individual. |
| $v_{ij}$ | Major variant of the $j$th infection of the $i$th individual |
| $\phi_i$ | Sex of the $i$th individual |
| $e_i$ | Age group of the $i$th individual |
| $f_i$ | Vaccine product (AstraZeneca or Pfizer) of the first vaccine dose of the $i$th individual |
| $h_i^r$ | Date of the $r$th vaccine dose of the $i$th individual |

**Table 3.**
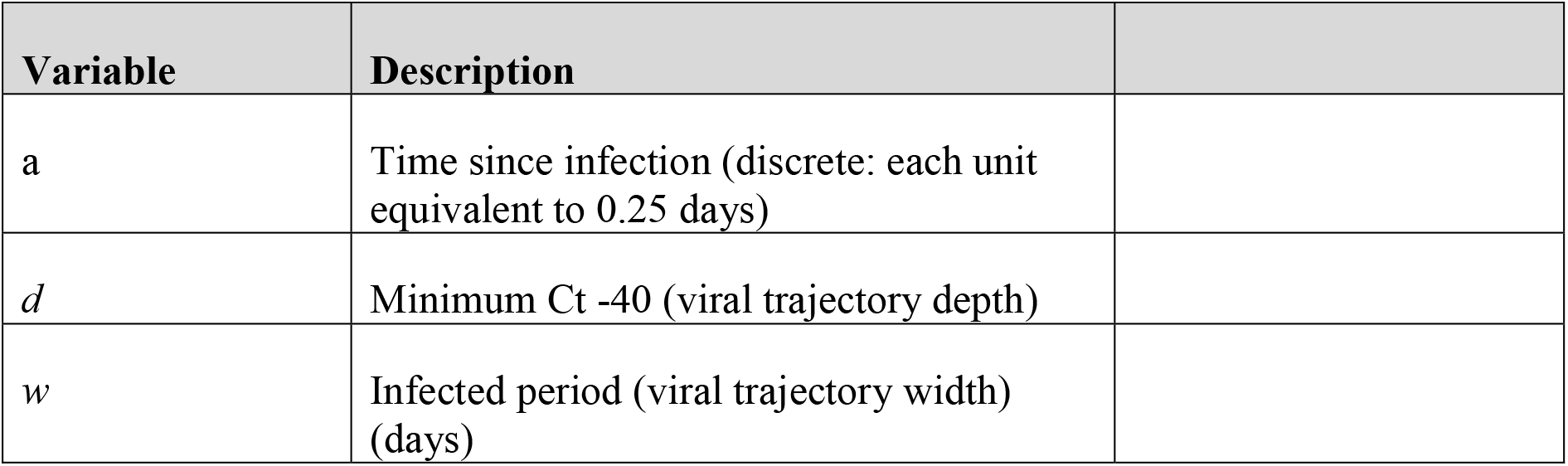
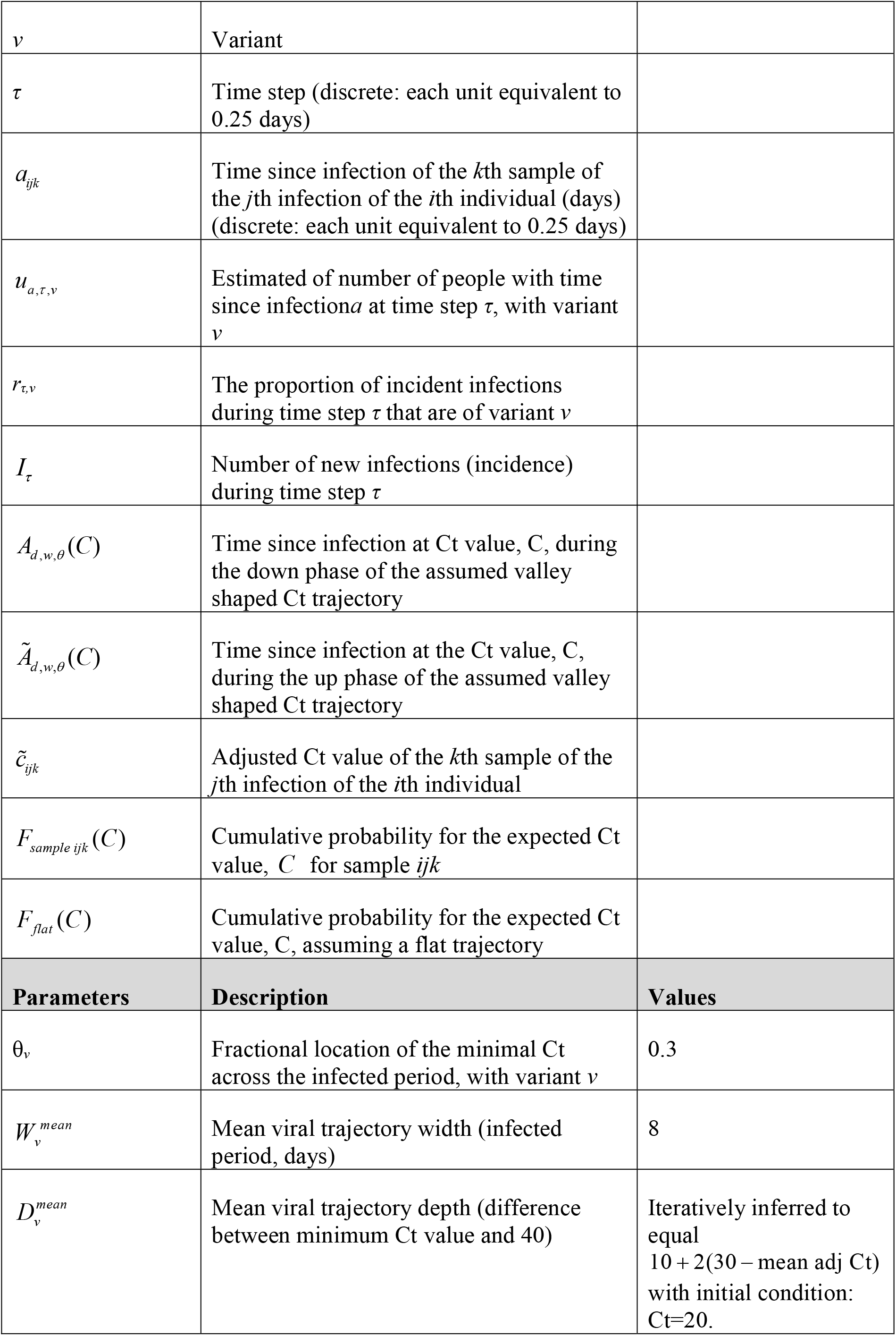
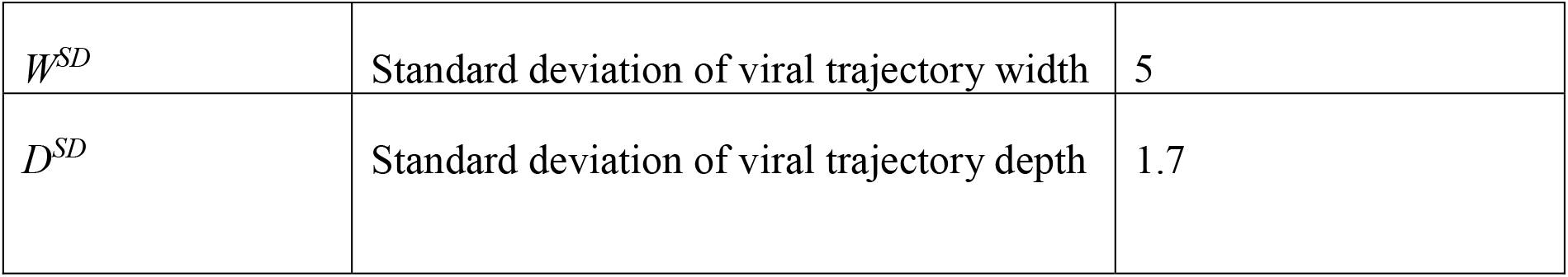
Description of additional variables and parameters used in calculation of adjusted Ct values.

| Variable | Description |
| --- | --- |
| a | Time since infection (discrete: each unit equivalent to 0.25 days) |
| d | Minimum Ct -40 (viral trajectory depth) |
| w | Infected period (viral trajectory width) (days) |

|  |  |  |
| --- | --- | --- |
| $v$ | Variant | |
| $\tau$ | Time step (discrete: each unit equivalent to 0.25 days) | |
| $a_{ijk}$ | Time since infection of the $k$th sample of the $j$th infection of the $i$th individual (days) (discrete: each unit equivalent to 0.25 days) | |
| $u_{a,\tau,v}$ | Estimated of number of people with time since infection $a$ at time step $\tau$, with variant $v$ | |
| $r_{\tau,v}$ | The proportion of incident infections during time step $\tau$ that are of variant $v$ | |
| $I_{\tau}$ | Number of new infections (incidence) during time step $\tau$ | |
| $A_{d,w,\theta}(C)$ | Time since infection at Ct value, $C$, during the down phase of the assumed valley shaped Ct trajectory | |
| $\tilde{A}_{d,w,\theta}(C)$ | Time since infection at the Ct value, $C$, during the up phase of the assumed valley shaped Ct trajectory | |
| $\tilde{c}_{ijk}$ | Adjusted Ct value of the $k$th sample of the $j$th infection of the $i$th individual | |
| $F_{sample\ ijk}(C)$ | Cumulative probability for the expected Ct value, $C$ for sample $ijk$ | |
| $F_{flat}(C)$ | Cumulative probability for the expected Ct value, $C$, assuming a flat trajectory | |
| **Parameters** | **Description** | **Values** |
| $\theta_v$ | Fractional location of the minimal Ct across the infected period, with variant $v$ | 0.3 |
| $W_v^{mean}$ | Mean viral trajectory width (infected period, days) | 8 |
| $D_v^{mean}$ | Mean viral trajectory depth (difference between minimum Ct value and 40) | Iteratively inferred to equal $10 + 2(30 - \text{mean adj Ct})$ with initial condition: Ct=20. |

|  |  |  |
| --- | --- | --- |
| $W^{SD}$ | Standard deviation of viral trajectory width | 5 |
| $D^{SD}$ | Standard deviation of viral trajectory depth | 1.7 |

Because several of the VIF values for other predictor variables among both sample sets were greater than 3, we analysed our data using PLS regression to acknowledge the difficulties in disentangling the relative roles of different factors in explaining viral burdens.

### Viral burdens are higher among older individuals

For samples sequenced in Oxford, six components (linear combinations of the predictors that are orthogonal to each other) describe the data (Supplementary Figure 1a), as determined by the number that minimises the mean squared prediction error. Although these components only explain a small amount of variance in the adjusted Ct values (2.1%), the first two are both significant in predicting the values in a quantile median regression model (p<0.0001 and p=0.003) (used to acknowledge non-normality in the residuals). For the Northumbria samples, six latent components also minimise the mean squared prediction error, the first three of which significantly predict (p<0.0001) the adjusted Ct values (Supplementary Figure 1a and 1b). This analysis highlights that, taken together, factors included in our model significantly impact viral burdens. For reference, loading plots for the first two latent components of each sample are shown in Supplementary Figures 1c and 1d.

Beta scores (which can be considered equivalent to regression coefficients), and variance in projection (VIP) scores can be used to assess the magnitude and importance of the contribution of the different variables to the response (Table 1), respectively. Variables with VIP values greater than 1 are typically considered to be important and those with VIP values greater than 0.8 are considered to be borderline important. Using this approach, we identified age as an important predictor of Ct values among both the Oxford (Beta score=-0.013 per year, VIP=1.38) and Northumbria (Beta score=-0.026 per year, VIP=2.15) samples. The effect that we measure equates to the mean Ct value being on average (across both datasets) 0.20 lower for every 10 years older.

There was no evidence of an association between sex and viral burden among either the Oxford (Beta score=-0.13, VIP=0.35) or Northumbria samples (Beta score=-0.13, VIP=0.63), with only slightly lower Ct values (higher viral burdens) in males compared to females.

### Among individuals with no known exposure, viral burdens are higher during Alpha compared to B.1.177 infection

We defined unvaccinated individuals with no known prior exposure as those individuals who have had neither a previous recorded infection, a previous positive test for spike antibodies, nor a vaccine at least 14 days prior. For samples sequenced at Oxford, Ct values in this group were higher for B.1.177 samples (Beta score=0.94, VIP=1.42) compared to Alpha. Since this difference in Ct values is in the opposite direction to that expected from increasing immunity over time, it is credible to infer that infection by the Alpha variant directly resulted in higher within-host viral burdens compared to infection with B.1.177.

Among unvaccinated individuals with no known prior exposure sampled at Oxford, we also found strong importance in support of Ct values being higher in Delta infected individuals (Beta score=1.06, VIP=1.52) than Alpha infected individuals. However, it is not possible to determine whether this difference is caused by infection by the different variants, or other factors that also correlate with calendar time. In particular, infection-acquired immunity has been increasing in the population over time, and we cannot rule out increased immunity over time, rather than the shift from the Alpha variant to the Delta variant, explaining the measured difference.

Among individuals with no known prior exposure whose samples were sequenced at Northumbria, there was no evidence of a difference in Ct values between BA.1 Omicron and Delta (Beta score=-0.14, VIP=0.60).

### Among vaccinated individuals, viral burdens are higher during Delta compared to Alpha infection

For individuals who were vaccinated or had a known prior exposure, we further categorised them according to whether they had either tested positive for spike antibodies prior to the first PCR-positive sample in the infection, or had 1 vaccine dose, 2 vaccine doses, or 3 vaccine doses. Individuals who had both a prior antibody positive sample and were vaccinated were assigned to the appropriate vaccination group (1, 2 or 3 doses). Among the Oxford samples, Ct values were higher among vaccinated individuals, compared to those with no known prior exposure (Beta score=1.42, VIP=1.33). Though the magnitude and importance of the signal was weaker, a similar pattern was observed among the Northumbria samples (Beta score=0.47, VIP=0.88). The impact of two vaccine doses over one on Ct values was limited (Oxford samples: Beta score=0.01, VIP=0.75; Northumbria samples: Beta score=0.05, VIP=0.82), but the impact of variant among vaccinated individuals was important. Ct values were lower among Delta compared to Alpha infections (Oxford samples: Beta score=-0.34, VIP=1.02) and, although less significant, also lower among BA.1 Omicron infections, compared to Delta infections (Northumbria samples: Beta score=-0.15, VIP=0.88). Vaccination with AstraZeneca was associated with slightly higher viral burdens compared to Pfizer (Oxford samples: Beta score=-0.09, VIP=1.02; Northumbria samples: Beta score=-0.09, VIP=1.00).

There was an effect of lower higher Ct values among individuals with a prior antibody positive sample (compared to those with no known prior exposure); however, the importance of this factor was low (Oxford samples: Beta score=3.29, VIP=0.42; Northumbria samples: Beta score=0.33, VIP=0.24), as was the impact of variant in this group.

### Higher viral burden in Alpha infections is robust to assumptions about the within-host viral trajectory

Given our previous observation that the assumed asymmetry in the viral load trajectory can have a measurable impact on the adjusted Ct value, we conducted a sensitivity analysis on our PLS regression. We varied the parameter that determines the asymmetry of the within-host viral burden trajectory for each of the variants. Both the Beta score (Figure 5a) and VIP value (Figure 5b) the for the indicator for the variant being B.1.177 rather than Alpha among individuals sampled at Oxford with no known prior exposure (i.e. no known prior infection, prior antibodies or vaccination) decreased as the assumed viral burden trajectories of B.1.177 were more skewed towards the start of the infection compared to Alpha (Figures 5a and 5b). These relationships are linked to the fact that although the Oxford samples span the whole of the Alpha wave, they did not span the early part of the B.1.177 wave. It is noteworthy that the VIP value remained greater than unity across plausible parameter combinations, providing support for the conclusion that viral burdens are higher in samples with the Alpha variant compared to B.1.177, among these individuals.

**Figure 5.**
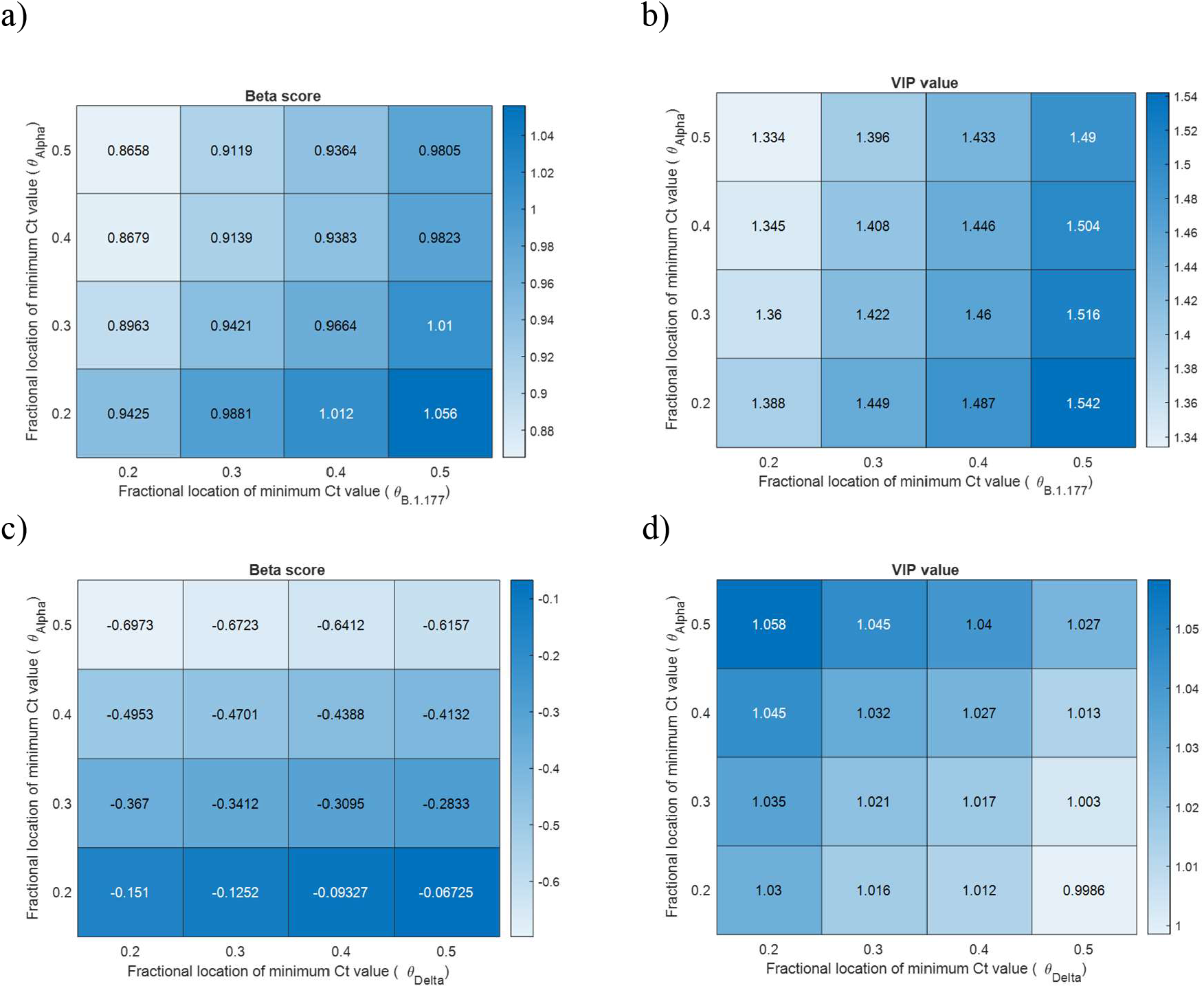
Sensitivity analysis investigating the impact of the shape of within-host viral trajectory on PLS regression analysis into the impact of variant on Ct values. Panels a) and c) show Beta scores, which can be considered to be equivalent to regression coefficients, defining the magnitude of the effect of the variant on the adjusted Ct values. Panels b) and d) show VIP values defining the importance of the association – where values greater than 1 are typically considered to indicate importance. Panels a) and b) investigate the association between the variant being B.1.177 (relative to Alpha) and Ct values among individuals with no known prior immunity. The Beta scores and VIP values vary with changes to the assumed asymmetry of the within-host viral burden trajectory associated with the B.1.177 lineage and the Alpha variant. The asymmetry is determined by changes to the fractional location of the minimum Ct value (peak viral burden) for each variant (*θ*_B.1.177_ and *θ*_Alpha_, respectively). Data sampled at Oxford. Panels c) and d) investigate the association between the variant being Delta (relative to Alpha) and Ct values among vaccinated individuals and how the Beta scores and VIP values vary with changes to *θ*_Alpha_ and *θ*_Delta_, respectively. Data sampled at Oxford.

When evaluating the impact on Ct values of the variant being Delta (rather than Alpha) among vaccinated individuals (Figures 5c and 5d), both the VIP value and the magnitude of the Beta score increased as the assumed viral burdens of Delta were more skewed towards the start of the infection compared to Alpha. These relationships are linked to the fact that that the Oxford samples did not span the latter part of the Delta wave. The VIP value remained greater than unity (or very close to for higher discordant within-host trajectories of the two variants) across plausible parameter combinations. This analysis therefore provides support for the finding that samples with the Delta variant had lower viral burdens compared to samples with the Alpha variant among vaccinated individuals.

## Discussion

We developed a framework to compare within-host viral burdens across different SARS-CoV-2 variants from random survey data, such as the CIS. The method directly estimates the level of uncertainty in the time-since-infection of each sample due to the sparse nature of the sampling and the effect of differing epidemiological trends of SARS-CoV-2 variants. The method highlights how the combination of the within-host viral trajectory and the epidemiological trajectory of a viral variant can influence observed viral burdens in survey data.

Using this framework, we inferred epidemiologically adjusted Ct values from samples sequenced as part of the CIS, a large-scale community survey, recruiting randomly selected private residential households and testing participants regardless of symptoms. Using partial least squares regression we found that viral burdens were higher among older individuals. In addition, among individuals with no known prior immunity, viral burdens were, on average, higher among Alpha-variant compared to B.1.177 samples. Viral burdens among individuals with no known prior exposure to infection or vaccination then decreased during the transition from primarily Alpha to primarily Delta infections. However, it is not possible to determine whether this was due to the infecting variant or other factors that also have a temporal component. For example, an increase in immunity due to unobserved infection over time could also explain this result. Among vaccinated individuals, we found evidence of higher viral burdens in infections with the Delta-variant, compared to those with the Alpha-variant.

Our study supports the hypothesis that the observed increases in transmissibility from B.1.177 to the Alpha variant (in individuals with no known prior exposure to infection or vaccination) and then to the Delta variant (in vaccinated individuals) were, at least in part, due to higher viral burdens. However, we cannot rule out other factors playing a role, including differences between variants in the viral shedding rate, infectious period [25], or per-virion probability of transmission. Although we infer higher viral burdens among BA.1 Omicron samples relative to Delta variant samples, our inferred support for this result is not strong. The replacement of the Delta variant with the BA.1 Omicron variant in the UK therefore cannot be clearly attributed to changes in viral burdens.

For this study we determined viral variant from viral sequence data, which in practice meant excluding samples with low viral burdens. This is because only samples with Ct ≤30 are routinely sequenced, and additionally, samples with higher Ct values (lower viral burdens) are less likely to have sufficient genomic coverage to determine the variant. Although these restrictions could impact our qualitative estimates, we do not expect them to bias our main qualitative results. Furthermore, since individuals with low viral burdens contribute little to viral transmission [26], our study reflects the impact of viral variants and other factors on viral burdens at levels that are relevant for transmission.

Monitoring of the characteristics of SARS-CoV-2 variants will continue to be critical to public health decisions in the foreseeable future. As more countries roll out population representative surveys, correcting for epidemiological effects will remain important. More generally, any studies using community surveillance data that aim to consider traits that vary through infection (e.g. Ct values, immune markers), could be impacted by pathogen epidemiology and therefore could benefit from epidemiological adjustment. In summary, our study promotes a new way of critically analysing random survey data to acknowledge the combined impact of pathogen epidemiology and within-host traits that vary over the course of an infection.

## Methods

### Study cohort

We used data from the Office for National Statistics Covid infection survey (ISRCTN21086382CT, https://www.ndm.ox.ac.uk/covid-19/covid-19-infection-survey). The survey has been described in detail elsewhere [24]. However, in brief, private households were randomly selected on a continuing basis in order to provide a representative sample of inhabitants of the UK. Following agreement to participate, self-collected nose and throat swabs were taken by participants – or their parents/carers if under 12 years of age – as instructed by a study worker. The intended schedule of swabbing was weekly for the first month of participation and monthly thereafter, for up to a year. However, there was variability among participants due to missed or late swabs, and participants could also chose to participate only once, or only for the first month, rather than on an ongoing basis, and were also free to leave the study at any time. For a random 10–20% of households, participants 16 years or older were invited to provide monthly venous blood samples for assays of anti-trimeric spike protein IgG. Metadata that includes age, sex, gender, postcode and vaccination details, were additionally recorded.

### Sequencing and lineage identification

All swabs were tested for SARS-CoV-2 using RT-QPCR, and the cycle threshold (Ct) values of positive samples were recorded. A random selection of positive samples collected before mid-December 2020 were sequenced, and from mid-December 2020 onwards the ambition was to sequence all positive samples with Ct≤30. Sequenced samples collected between 27^th^ Sep 2020 and 17^th^ July 2021 were sequenced at the University of Oxford using veSEQ. This employs an RNASeq protocol based on a quantitative targeted enrichment strategy [27] and sequencing on the Illumina Novaseq platform. For a full description of the sequencing protocol see [27, 28]. Most sequenced samples collected between 20^th^ Sep 2021 and 19^th^ Jan 2022 were sequenced at the University of Northumbria using the CoronaHiT [29] variant of the ARTIC protocol and Illumina Novaseq 550. Consensus sequences were produced using the *shiver* pipeline [30] and lineage assigned using the PangoLEARN [31].

All samples sequenced in Oxford with Ct≤30 were retained for analysis, with the added restriction of ≥50% genome coverage required for samples sequenced in Northumbria. Lineages were assigned using the PangoLEARN [31], with samples assigned as B.1.177 (and sublineages), Alpha (B.1.1.7 and sublineages), Delta (B.1.617.2 and sublineages) and Omicron (BA.1 and sublineages) used for this analysis. For Oxford sequenced samples with <50% coverage, and which could not be reliably assigned using PangoLEARN, we assigned one of the four major lineages if a consensus base was called at three or more lineage defining sites, and with more than two-thirds of these calls consistent with the lineage. To avoid differences in sequencing protocol influencing our analyses, samples sequenced in Oxford and Northumbria were analysed separately.

### Infection characteristics

All individuals with at least one positive sample sequenced in Oxford or Northumbria, and with the virus assigned to one of the four major lineages as described above, were included in our analysis, and indexed *i=1*…*n*, where *n* is the number of individuals. If an individual was infected by more than one major lineage during the study period, these were designated with an infection number *j*, where *j=1* represents the first infection, *j=2* the second infection, and so on. Positive samples were assumed to be part of the same infection if they were of the same major variant and were in a continuous sequence of positive samples (i.e. no negative intermediate samples). The index *k* denotes the *k*th sample of the infection. In the case of a non-continuous sequence of positive samples of the same major lineage, any addition positive samples were excluded from our study. Infections which were of the same major lineage but not in a continuous sequence of positive samples were excluded from the analysis. The list of variables used to describe the data are given in Table 2.

### Calculating epidemiologically adjusted Ct values

#### Step 1. Describing the within-host Ct trajectory

We assume that within-host Ct trajectories are piecewise linear and valley-shaped (Figure 1b), defined by the infected period (width, *w*) and the difference between the minimum Ct value and 40 (depth, *d*). Probability distributions for these variables (calculated in a discrete manner, each spaced by value 0.25 and 0.5 respectively) are derived from truncated discretised normal distributions, described by *p(d)* (equation 1) and *p(w)* (equation 2), with means 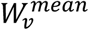 *and* 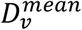 and standard deviations, *W*^*SD*^, *D*^*SD*^, so that

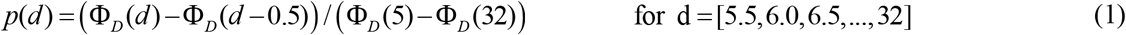

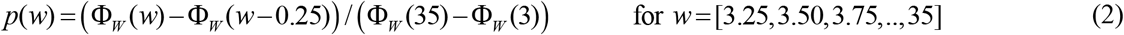

where

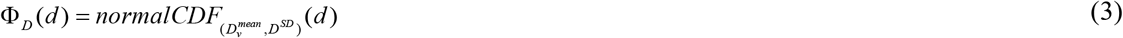

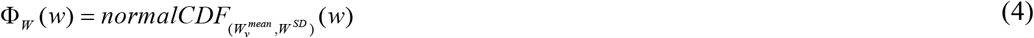

The peak viral burden is assumed to occur at a time since infection equal to a fraction, θ<1, of the total infected period. The parameters 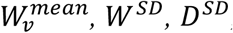, and θ are derived from previous studies and varied in sensitivity analyses. The parameter 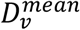, is iteratively inferred to a tolerance of 0.1 following implementation of the methodology described – which, for each sample, estimates an adjusted Ct value – and calculated to equal twice the difference between 40 and the mean adjusted Ct value for that variant. For ease of reference, all other variables described here and throughout the following derivation are listed in Table 3.

#### Step 2. Estimating the distribution of time since infection for different SARS-CoV-2 variants over calendar time

We estimated the distribution of infections in the population stratified by variant and time since infection over calendar time using published estimates of total incidence of SARS-CoV-2 in the UK (www.ons.gov.uk/peoplepopulationandcommunity/healthandsocialcare/conditionsanddiseases/datasets/coronaviruscovid19infectionsurveydata) and published estimates of the proportion of incident infections with each of the major variants under study (B.1.177, Alpha, Delta and BA.1 Omicron) over time from the COVID-19 Genomics UK Consortium (COG-UK: www.cogconsortium.uk). Working in discrete time steps (τ=1,2,3…) that are 0.25 days each, we define *I*_*τ*_ to be the incidence during time step, *τ* and *r*_*τ,v*_ to be the proportion of incident infections during time step *τ* that are of variant *v* (*v*=1:4 represent B.1.177, Alpha, Delta and BA.1 Omicron, respectively). We further define *u*_*a,τ,v*_ to be the number of infections with time since infection, *a* (stratified as discrete time steps of 0.25 days each), during time step *τ* with variant *v*. The number of incident infections (i.e. infections with time since infection=0) during time step *τ* with each variant *v* is estimated to be the product of the total incidence during that time step and the fraction of incident infections of that variant (*u*_0,*τ,v*_ = *r*_*τ,v*_ *I*_*τ*_). To estimate *u*_*a,τ,v*_ for each *a*>*0*, we assume that the infected periods are taken from a truncated normal distribution with mean, 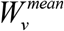, and variance *W*^*SD*^. Therefore, the number of infections of time since infection *a*, at time step *τ* is calculated to be the number of incident infections from time step *τ-a* that are still persisting after a time *a*, thus: 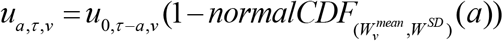.

#### Step 3. For each sample and each infected period, estimate a time since infection distribution

For each sample and for each assumed infected period (*w*), we inferred the distribution of time since infection. We first selected the distribution (Step 2) that corresponds to the sample date and variant of the sample and adjusted it to account for known bounds on the time since infection for that sample, measured in days. The bounds 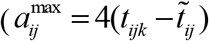 and 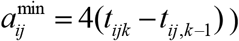 are derived by considering information on Ct values at previous samples and scaled to account for the transformation to discrete time steps. The time since infection probability distribution for each sample is then given by:

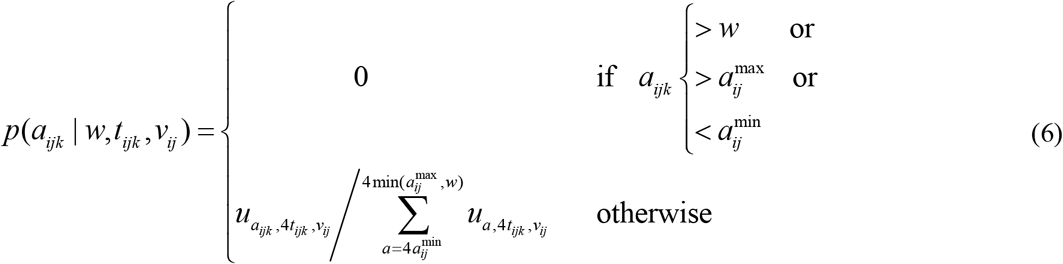

#### Step 4. Infer a sample-specific expected distribution of Ct values

For each sample, based upon the sample time (*t*_*ijk*_) and variant (*v*_*ij*_), we derived an expected distribution of Ct values (equation 7). This was done by conditioning on the time since infection (*a*) and the depth (*d*) and width (*w*) of the within host viral trajectory. These conditional probabilities were combined with the time since infection distributions derived in step 3 and the within-host parameter distributions described in step 1.

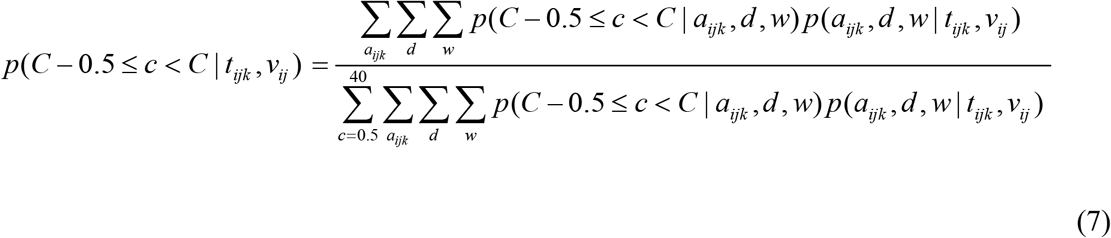

where the probability of a particular time since infection (*a*^*ijk*^), trajectory width (*w*) and trajectory depth (*d*) is given by:

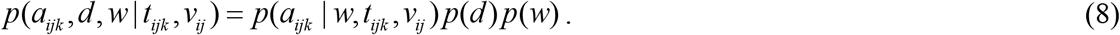

and the probability of the Ct value (*c*) falling within a certain discrete boundary, given the time since infection and the width and depth of the viral trajectory, is defined as 1 or 0 depending upon whether it matches up with the valley-shaped viral trajectory curve (figure 1b), as shown below:

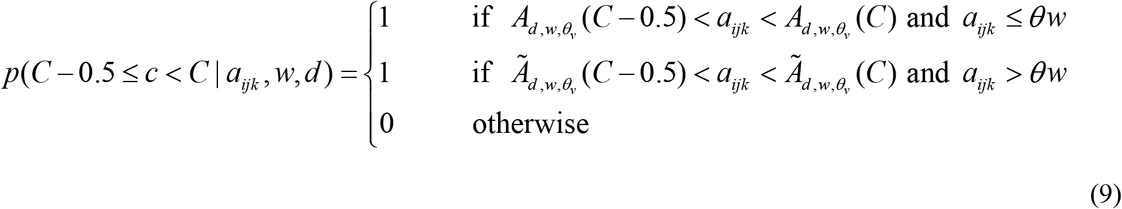

Where C is a dummy variable representing the Ct value, and

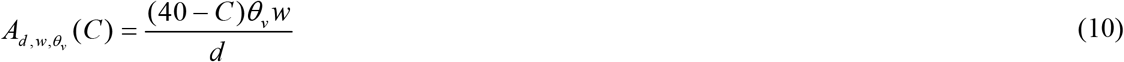

and

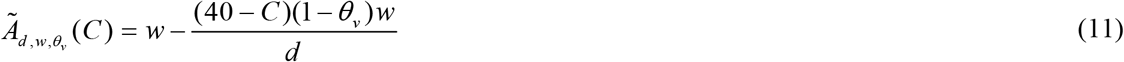

are dummy variables that describe the relationship between the Ct value (C) and the time since infection 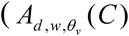 and 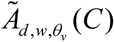, during down phase and up phase of the valley-shaped trajectory, respectively.

#### Step 5. Calculate an expected distribution of Ct values for a flat epidemic trajectory

The full process for calculating an expected distribution of Ct values (steps 1-4) was repeated under an assumption of a flat epidemic trajectory, rather than a variant-specific trajectory.

#### Step 6. For each sample, infer an epidemiologically adjusted Ct value

For each sample, we identified the percentile that the observed Ct (*c*_*ijk*_) falls in, among the sample-specific expected Ct distribution. The adjusted Ct value 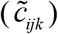 was then derived by identifying the Ct value at that percentile within the expected distribution of Ct values based upon a flat epidemic trajectory (Figure 1c).

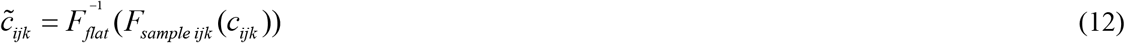

Where

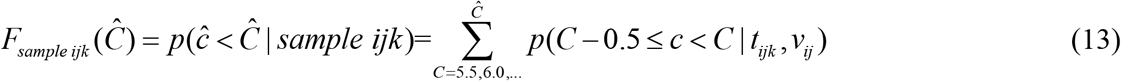

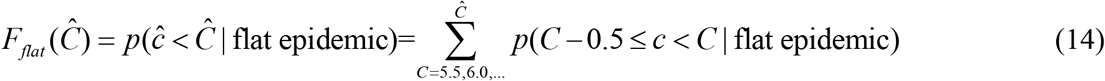

### Implementation of analysis

All analyses were implemented in Matlab and the code is available at https://github.com/helenfryer1000000/epidemiologically-adjusted-viral-load. Estimation of adjusted Ct values was implemented using a bespoke script. Partial least squares regression was implemented using the PLSregress function, which is part of the Statistics and Machine Learning toolbox in Matlab. Quantile median regression was implemented using the function qr_standard, provided at: https://github.com/zjph602xtc/Quantile_reg.

## Acknowledgements

K.A.L. and H.F. were supported by The Wellcome Trust and The Royal Society (107652/Z/15/Z to K.A.L..) and by the Li Ka Shing Foundation funding awarded to K.A.L.

COG-UK is supported by funding from the Medical Research Council (MRC) part of UK Research & Innovation (UKRI), the National Institute of Health Research (NIHR) [grant code: MC_PC_19027], and Genome Research Limited, operating as the Wellcome Sanger Institute. The authors acknowledge use of data generated through the COVID-19 Genomics Programme funded by the Department of Health and Social Care. The views expressed are those of the author and not necessarily those of the Department of Health and Social Care or UKHSA

LP gratefully acknowledges funding from the Wellcome Trust and Royal Society (grant number 202562/Z/16/Z), the UKRI through the JUNIPER modelling consortium (grant number MR/V038613/1) and the Alan Turing Institute under the EPSRC grant (EP/N510129/1).

Mohammad Adnan Tariq. Contract research at Northumbria University funded by UKHSA as a DNA sequencing resilience site.

Darren Smith funded by UKHSA, COG-UK, Research England E3: HBBE and Northumbria University.

Andrew Nelson & Clare McCann,funded by UKHSA, and Northumbria University

Matt Bashton, funded by UKHSA, Research England E3:HBBE

Greg Young, funded by Research England E3:HBBE

Amy Trebes is funded by the Wellcome (grant reference 203141/Z/16/Z).

David Buck is funded by the Wellcome (grant reference 203141/Z/16/Z).

Paolo Piazza is funded by the Wellcome (grant reference 203141/Z/16/Z).

Lorne Lonie is funded by the Wellcome (grant reference 203141/Z/16/Z).

Angie Green is funded by the Wellcome (grant reference 203141/Z/16/Z).

---

**The COVID-19 Genomics UK (COG-UK) consortium**

**June 2021 V.3**

**Funding acquisition, Leadership and supervision, Metadata curation, Project administration, Samples and logistics, Sequencing and analysis, Software and analysis tools, and Visualisation:**

Dr Samuel C Robson PhD ^13, 84^

**Funding acquisition, Leadership and supervision, Metadata curation, Project administration, Samples and logistics, Sequencing and analysis, and Software and analysis tools:**

Dr Thomas R Connor PhD ^11, 74^ and Prof Nicholas J Loman PhD ^43^

**Leadership and supervision, Metadata curation, Project administration, Samples and logistics, Sequencing and analysis, Software and analysis tools, and Visualisation:**

Dr Tanya Golubchik PhD ^5^

**Funding acquisition, Leadership and supervision, Metadata curation, Samples and logistics, Sequencing and analysis, and Visualisation:**

Dr Rocio T Martinez Nunez PhD ^46^

**Funding acquisition, Leadership and supervision, Project administration, Samples and logistics, Sequencing and analysis, and Software and analysis tools:**

Dr David Bonsall PhD ^5^

**Funding acquisition, Leadership and supervision, Project administration, Sequencing and analysis, Software and analysis tools, and Visualisation:**

Prof Andrew Rambaut DPhil ^104^

**Funding acquisition, Metadata curation, Project administration, Samples and logistics, Sequencing and analysis, and Software and analysis tools:**

Dr Luke B Snell MSc, MBBS ^12^

**Leadership and supervision, Metadata curation, Project administration, Samples and logistics, Software and analysis tools, and Visualisation:**

Rich Livett MSc ^116^

**Funding acquisition, Leadership and supervision, Metadata curation, Project administration, and Samples and logistics:**

Dr Catherine Ludden PhD ^20, 70^

**Funding acquisition, Leadership and supervision, Metadata curation, Samples and logistics, and Sequencing and analysis:**

Dr Sally Corden PhD ^74^ and Dr Eleni Nastouli FRCPath ^96, 95, 30^

**Funding acquisition, Leadership and supervision, Metadata curation, Sequencing and analysis, and Software and analysis tools:**

Dr Gaia Nebbia PhD, FRCPath ^12^

**Funding acquisition, Leadership and supervision, Project administration, Samples and logistics, and Sequencing and analysis:**

Ian Johnston BSc ^116^

**Leadership and supervision, Metadata curation, Project administration, Samples and logistics, and Sequencing and analysis:**

Prof Katrina Lythgoe PhD ^5^, Dr M. Estee Torok FRCP ^19, 20^ and Prof Ian G Goodfellow PhD ^24^

**Leadership and supervision, Metadata curation, Project administration, Samples and logistics, and Visualisation:**

Dr Jacqui A Prieto PhD ^97, 82^ and Dr Kordo Saeed MD, FRCPath ^97, 83^

**Leadership and supervision, Metadata curation, Project administration, Sequencing and analysis, and Software and analysis tools:**

Dr David K Jackson PhD ^116^

**Leadership and supervision, Metadata curation, Samples and logistics, Sequencing and analysis, and Visualisation:**

Dr Catherine Houlihan PhD ^96, 94^

**Leadership and supervision, Metadata curation, Sequencing and analysis, Software and analysis tools, and Visualisation:**

Dr Dan Frampton PhD ^94, 95^

**Metadata curation, Project administration, Samples and logistics, Sequencing and analysis, and Software and analysis tools:**

Dr William L Hamilton PhD ^19^ and Dr Adam A Witney PhD ^41^

**Funding acquisition, Samples and logistics, Sequencing and analysis, and Visualisation:**

Dr Giselda Bucca PhD ^101^

**Funding acquisition, Leadership and supervision, Metadata curation, and Project administration:**

Dr Cassie F Pope PhD^40, 41^

**Funding acquisition, Leadership and supervision, Metadata curation, and Samples and logistics:**

Dr Catherine Moore PhD ^74^

**Funding acquisition, Leadership and supervision, Metadata curation, and Sequencing and analysis:**

Prof Emma C Thomson PhD, FRCP ^53^

**Funding acquisition, Leadership and supervision, Project administration, and Samples and logistics:**

Dr Teresa Cutino-Moguel PhD ^2^, Dr Ewan M Harrison PhD ^116, 102^

**Funding acquisition, Leadership and supervision, Sequencing and analysis, and Visualisation:**

Prof Colin P Smith PhD ^101^

**Leadership and supervision, Metadata curation, Project administration, and Sequencing and analysis:**

Fiona Rogan BSc ^77^

**Leadership and supervision, Metadata curation, Project administration, and Samples and logistics:**

Shaun M Beckwith MSc ^6^, Abigail Murray Degree ^6^, Dawn Singleton HNC ^6^, Dr Kirstine Eastick PhD, FRCPath ^37^, Dr Liz A Sheridan PhD ^98^, Paul Randell MSc, PgD ^99^, Dr Leigh M Jackson PhD ^105^, Dr Cristina V Ariani PhD ^116^ and Dr Sónia Gonçalves PhD ^116^

**Leadership and supervision, Metadata curation, Samples and logistics, and Sequencing and analysis:**

Dr Derek J Fairley PhD ^3, 77^, Prof Matthew W Loose PhD ^18^ and Joanne Watkins MSc ^74^

**Leadership and supervision, Metadata curation, Samples and logistics, and Visualisation:**

Dr Samuel Moses MD ^25, 106^

**Leadership and supervision, Metadata curation, Sequencing and analysis, and Software and analysis tools:**

Dr Sam Nicholls PhD ^43^, Dr Matthew Bull PhD ^74^ and Dr Roberto Amato PhD ^116^

**Leadership and supervision, Project administration, Samples and logistics, and Sequencing and analysis:**

Prof Darren L Smith PhD ^36, 65, 66^

**Leadership and supervision, Sequencing and analysis, Software and analysis tools, and Visualisation:**

Prof David M Aanensen PhD ^14, 116^ and Dr Jeffrey C Barrett PhD ^116^

**Metadata curation, Project administration, Samples and logistics, and Sequencing and analysis:**

Dr Beatrix Kele PhD ^2^, Dr Dinesh Aggarwal MRCP^20, 116, 70^, Dr James G Shepherd MBCHB, MRCP ^53^, Dr Martin D Curran PhD ^71^ and Dr Surendra Parmar PhD ^71^

**Metadata curation, Project administration, Sequencing and analysis, and Software and analysis tools:**

Dr Matthew D Parker PhD ^109^

**Metadata curation, Samples and logistics, Sequencing and analysis, and Software and analysis tools:**

Dr Catryn Williams PhD ^74^

**Metadata curation, Samples and logistics, Sequencing and analysis, and Visualisation:**

Dr Sharon Glaysher PhD ^68^

**Metadata curation, Sequencing and analysis, Software and analysis tools, and Visualisation:**

Dr Anthony P Underwood PhD ^14, 116^, Dr Matthew Bashton PhD ^36, 65^, Dr Nicole Pacchiarini PhD ^74^, Dr Katie F Loveson PhD ^84^ and Matthew Byott MSc ^95, 96^

**Project administration, Sequencing and analysis, Software and analysis tools, and Visualisation:**

Dr Alessandro M Carabelli PhD ^20^

**Funding acquisition, Leadership and supervision, and Metadata curation:**

Dr Kate E Templeton PhD ^56, 104^

**Funding acquisition, Leadership and supervision, and Project administration:**

Dr Thushan I de Silva PhD ^109^, Dr Dennis Wang PhD ^109^, Dr Cordelia F Langford PhD ^116^ and John Sillitoe BEng ^116^

**Funding acquisition, Leadership and supervision, and Samples and logistics:**

Prof Rory N Gunson PhD, FRCPath ^55^

**Funding acquisition, Leadership and supervision, and Sequencing and analysis:**

Dr Simon Cottrell PhD ^74^, Dr Justin O’Grady PhD ^75, 103^ and Prof Dominic Kwiatkowski PhD ^116, 108^

**Leadership and supervision, Metadata curation, and Project administration:**

Dr Patrick J Lillie PhD, FRCP ^37^

**Leadership and supervision, Metadata curation, and Samples and logistics:**

Dr Nicholas Cortes MBCHB ^33^, Dr Nathan Moore MBCHB ^33^, Dr Claire Thomas DPhil ^33^, Phillipa J Burns MSc, DipRCPath ^37^, Dr Tabitha W Mahungu FRCPath ^80^ and Steven Liggett BSc ^86^

**Leadership and supervision, Metadata curation, and Sequencing and analysis:**

Angela H Beckett MSc ^13, 81^ and Prof Matthew TG Holden PhD ^73^

**Leadership and supervision, Project administration, and Samples and logistics:**

Dr Lisa J Levett PhD ^34^, Dr Husam Osman PhD ^70, 35^ and Dr Mohammed O Hassan-Ibrahim PhD, FRCPath ^99^

**Leadership and supervision, Project administration, and Sequencing and analysis:**

Dr David A Simpson PhD ^77^

**Leadership and supervision, Samples and logistics, and Sequencing and analysis:**

Dr Meera Chand PhD ^72^, Prof Ravi K Gupta PhD ^102^, Prof Alistair C Darby PhD ^107^ and Prof Steve Paterson PhD ^107^

**Leadership and supervision, Sequencing and analysis, and Software and analysis tools:**

Prof Oliver G Pybus DPhil ^23^, Dr Erik M Volz PhD ^39^, Prof Daniela de Angelis PhD ^52^, Prof David L Robertson PhD ^53^, Dr Andrew J Page PhD ^75^ and Dr Inigo Martincorena PhD ^116^

**Leadership and supervision, Sequencing and analysis, and Visualisation:**

Dr Louise Aigrain PhD ^116^ and Dr Andrew R Bassett PhD ^116^

**Metadata curation, Project administration, and Samples and logistics:**

Dr Nick Wong DPhil, MRCP, FRCPath ^50^, Dr Yusri Taha MD, PhD ^89^, Michelle J Erkiert BA ^99^ and Dr Michael H Spencer Chapman MBBS ^116, 102^

**Metadata curation, Project administration, and Sequencing and analysis:**

Dr Rebecca Dewar PhD ^56^ and Martin P McHugh MSc ^56, 111^

**Metadata curation, Project administration, and Software and analysis tools:**

Siddharth Mookerjee MPH ^38, 57^

**Metadata curation, Project administration, and Visualisation:**

Stephen Aplin ^97^, Matthew Harvey ^97^, Thea Sass ^97^, Dr Helen Umpleby FRCP ^97^ and Helen Wheeler ^97^

**Metadata curation, Samples and logistics, and Sequencing and analysis:**

Dr James P McKenna PhD ^3^, Dr Ben Warne MRCP ^9^, Joshua F Taylor MSc ^22^, Yasmin Chaudhry BSc ^24^, Rhys Izuagbe ^24^, Dr Aminu S Jahun PhD ^24^, Dr Gregory R Young PhD ^36, 65^, Dr Claire McMurray PhD ^43^, Dr Clare M McCann PhD ^65, 66^, Dr Andrew Nelson PhD ^65, 66^ and Scott Elliott ^68^

**Metadata curation, Samples and logistics, and Visualisation:**

Hannah Lowe MSc ^25^

**Metadata curation, Sequencing and analysis, and Software and analysis tools:**

Dr Anna Price PhD ^11^, Matthew R Crown BSc ^65^, Dr Sara Rey PhD ^74^, Dr Sunando Roy PhD ^96^ and Dr Ben Temperton PhD ^105^

**Metadata curation, Sequencing and analysis, and Visualisation:**

Dr Sharif Shaaban PhD ^73^ and Dr Andrew R Hesketh PhD ^101^

**Project administration, Samples and logistics, and Sequencing and analysis:**

Dr Kenneth G Laing PhD^41^, Dr Irene M Monahan PhD ^41^ and Dr Judith Heaney PhD ^95, 96, 34^

**Project administration, Samples and logistics, and Visualisation:**

Dr Emanuela Pelosi FRCPath ^97^, Siona Silviera MSc ^97^ and Dr Eleri Wilson-Davies MD, FRCPath ^97^

**Samples and logistics, Software and analysis tools, and Visualisation:**

Dr Helen Fryer PhD ^5^

**Sequencing and analysis, Software and analysis tools, and Visualization:**

Dr Helen Adams PhD ^4^, Dr Louis du Plessis PhD ^23^, Dr Rob Johnson PhD ^39^, Dr William T Harvey PhD ^53, 42^, Dr Joseph Hughes PhD ^53^, Dr Richard J Orton PhD ^53^, Dr Lewis G Spurgin PhD ^59^, Dr Yann Bourgeois PhD ^81^, Dr Chris Ruis PhD ^102^, Áine O’Toole MSc ^104^, Marina Gourtovaia MSc ^116^ and Dr Theo Sanderson PhD ^116^

**Funding acquisition, and Leadership and supervision:**

Dr Christophe Fraser PhD ^5^, Dr Jonathan Edgeworth PhD, FRCPath ^12^, Prof Judith Breuer MD ^96, 29^, Dr Stephen L Michell PhD ^105^ and Prof John A Todd PhD ^115^

**Funding acquisition, and Project administration:**

Michaela John BSc ^10^ and Dr David Buck PhD ^115^

**Leadership and supervision, and Metadata curation:**

Dr Kavitha Gajee MBBS, FRCPath ^37^ and Dr Gemma L Kay PhD ^75^

**Leadership and supervision, and Project administration:**

Prof Sharon J Peacock PhD ^20, 70^ and David Heyburn ^74^

**Leadership and supervision, and Samples and logistics:**

Dr Themoula Charalampous PhD ^12, 46^, Adela Alcolea-Medina ^32, 112^, Katie Kitchman BSc ^37^, Prof Alan McNally PhD ^43, 93^, David T Pritchard MSc, CSci ^50^, Dr Samir Dervisevic FRCPath ^58^, Dr Peter Muir PhD ^70^, Dr Esther Robinson PhD ^70, 35^, Dr Barry B Vipond PhD ^70^, Newara A Ramadan MSc, CSci, FIBMS ^78^, Dr Christopher Jeanes MBBS ^90^, Danni Weldon BSc ^116^, Jana Catalan MSc ^118^ and Neil Jones MSc ^118^

**Leadership and supervision, and Sequencing and analysis:**

Dr Ana da Silva Filipe PhD ^53^, Dr Chris Williams MBBS ^74^, Marc Fuchs BSc ^77^, Dr Julia Miskelly PhD ^77^, Dr Aaron R Jeffries PhD ^105^, Karen Oliver BSc ^116^ and Dr Naomi R Park PhD ^116^

**Metadata curation, and Samples and logistics:**

Amy Ash BSc ^1^, Cherian Koshy MSc, CSci, FIBMS ^1^, Magdalena Barrow ^7^, Dr Sarah L Buchan PhD ^7^, Dr Anna Mantzouratou PhD ^7^, Dr Gemma Clark PhD ^15^, Dr Christopher W Holmes PhD ^16^, Sharon Campbell MSc ^17^, Thomas Davis MSc ^21^, Ngee Keong Tan MSc ^22^, Dr Julianne R Brown PhD ^29^, Dr Kathryn A Harris PhD ^29, 2^, Stephen P Kidd MSc ^33^, Dr Paul R Grant PhD ^34^, Dr Li Xu-McCrae PhD ^35^, Dr Alison Cox PhD ^38, 63^, Pinglawathee Madona ^38, 63^, Dr Marcus Pond PhD ^38, 63^, Dr Paul A Randell MBBCh ^38, 63^, Karen T Withell FIBMS ^48^, Cheryl Williams MSc ^51^, Dr Clive Graham MD ^60^, Rebecca Denton-Smith BSc ^62^, Emma Swindells BSc ^62^, Robyn Turnbull BSc ^62^, Dr Tim J Sloan PhD ^67^, Dr Andrew Bosworth PhD ^70, 35^, Stephanie Hutchings ^70^, Hannah M Pymont MSc ^70^, Dr Anna Casey PhD ^76^, Dr Liz Ratcliffe PhD ^76^, Dr Christopher R Jones PhD ^79, 105^, Dr Bridget A Knight PhD ^79, 105^, Dr Tanzina Haque PhD, FRCPath ^80^, Dr Jennifer Hart MRCP ^80^, Dr Dianne Irish-Tavares FRCPath ^80^, Eric Witele MSc ^80^, Craig Mower BA ^86^, Louisa K Watson DipHE ^86^, Jennifer Collins BSc ^89^, Gary Eltringham BSc ^89^, Dorian Crudgington ^98^, Ben Macklin ^98^, Prof Miren Iturriza-Gomara PhD ^107^, Dr Anita O Lucaci PhD ^107^ and Dr Patrick C McClure PhD ^113^

**Metadata curation, and Sequencing and analysis:**

Matthew Carlile BSc ^18^, Dr Nadine Holmes PhD ^18^, Dr Christopher Moore PhD ^18^, Dr Nathaniel Storey PhD ^29^, Dr Stefan Rooke PhD ^73^, Dr Gonzalo Yebra PhD ^73^, Dr Noel Craine DPhil ^74^, Malorie Perry MSc ^74^, Dr Nabil-Fareed Alikhan PhD ^75^, Dr Stephen Bridgett PhD ^77^, Kate F Cook MScR ^84^, Christopher Fearn MSc ^84^, Dr Salman Goudarzi PhD ^84^, Prof Ronan A Lyons MD ^88^, Dr Thomas Williams MD ^104^, Dr Sam T Haldenby PhD ^107^, Jillian Durham BSc ^116^ and Dr Steven Leonard PhD ^116^

**Metadata curation, and Software and analysis tools:**

Robert M Davies MA (Cantab) ^116^

**Project administration, and Samples and logistics:**

Dr Rahul Batra MD ^12^, Beth Blane BSc ^20^, Dr Moira J Spyer PhD ^30, 95, 96^, Perminder Smith MSc ^32, 112^, Mehmet Yavus ^85, 109^, Dr Rachel J Williams PhD ^96^, Dr Adhyana IK Mahanama MD ^97^, Dr Buddhini Samaraweera MD ^97^, Sophia T Girgis MSc ^102^, Samantha E Hansford CSci ^109^, Dr Angie Green PhD ^115^, Dr Charlotte Beaver PhD ^116^, Katherine L Bellis ^116, 102^, Matthew J Dorman ^116^, Sally Kay ^116^, Liam Prestwood ^116^ and Dr Shavanthi Rajatileka PhD ^116^

**Project administration, and Sequencing and analysis:**

Dr Joshua Quick PhD ^43^

**Project administration, and Software and analysis tools:**

Radoslaw Poplawski BSc ^43^

**Samples and logistics, and Sequencing and analysis:**

Dr Nicola Reynolds PhD ^8^, Andrew Mack MPhil ^11^, Dr Arthur Morriss PhD ^11^, Thomas Whalley BSc ^11^, Bindi Patel BSc ^12^, Dr Iliana Georgana PhD ^24^, Dr Myra Hosmillo PhD ^24^, Malte L Pinckert MPhil ^24^, Dr Joanne Stockton PhD ^43^, Dr John H Henderson PhD ^65^, Amy Hollis HND ^65^, Dr William Stanley PhD ^65^, Dr Wen C Yew PhD ^65^, Dr Richard Myers PhD ^72^, Dr Alicia Thornton PhD ^72^, Alexander Adams BSc ^74^, Tara Annett BSc ^74^, Dr Hibo Asad PhD ^74^, Alec Birchley MSc ^74^, Jason Coombes BSc ^74^, Johnathan M Evans MSc ^74^, Laia Fina ^74^, Bree Gatica-Wilcox MPhil ^74^, Lauren Gilbert ^74^, Lee Graham BSc ^74^, Jessica Hey BSc ^74^, Ember Hilvers MPH ^74^, Sophie Jones MSc ^74^, Hannah Jones ^74^, Sara Kumziene-Summerhayes MSc ^74^, Dr Caoimhe McKerr PhD ^74^, Jessica Powell BSc ^74^, Georgia Pugh ^74^, Sarah Taylor ^74^, Alexander J Trotter MRes ^75^, Charlotte A Williams BSc ^96^, Leanne M Kermack MSc ^102^, Benjamin H Foulkes MSc ^109^, Marta Gallis MSc ^109^, Hailey R Hornsby MSc ^109^, Stavroula F Louka MSc ^109^, Dr Manoj Pohare PhD ^109^, Paige Wolverson MSc ^109^, Peijun Zhang MSc ^109^, George MacIntyre-Cockett BSc ^115^, Amy Trebes MSc ^115^, Dr Robin J Moll PhD ^116^, Lynne Ferguson MSc ^117^, Dr Emily J Goldstein PhD ^117^, Dr Alasdair Maclean PhD ^117^ and Dr Rachael Tomb PhD ^117^

**Samples and logistics, and Software and analysis tools:**

Dr Igor Starinskij MSc, MRCP ^53^

**Sequencing and analysis, and Software and analysis tools:**

Laura Thomson BSc ^5^, Joel Southgate MSc ^11, 74^, Dr Moritz UG Kraemer DPhil ^23^, Dr Jayna Raghwani PhD ^23^, Dr Alex E Zarebski PhD ^23^, Olivia Boyd MSc ^39^, Lily Geidelberg MSc ^39^, Dr Chris J Illingworth PhD ^52^, Dr Chris Jackson PhD ^52^, Dr David Pascall PhD ^52^, Dr Sreenu Vattipally PhD ^53^, Timothy M Freeman MPhil ^109^, Dr Sharon N Hsu PhD ^109^, Dr Benjamin B Lindsey MRCP ^109^, Dr Keith James PhD ^116^, Kevin Lewis ^116^, Gerry Tonkin-Hill ^116^ and Dr Jaime M Tovar-Corona PhD ^116^

**Sequencing and analysis, and Visualisation:**

MacGregor Cox MSci ^20^

**Software and analysis tools, and Visualisation:**

Dr Khalil Abudahab PhD ^14, 116^, Mirko Menegazzo ^14^, Ben EW Taylor MEng ^14, 116^, Dr Corin A Yeats PhD ^14^, Afrida Mukaddas BTech ^53^, Derek W Wright MSc ^53^, Dr Leonardo de Oliveira Martins PhD ^75^, Dr Rachel Colquhoun DPhil ^104^, Verity Hill ^104^, Dr Ben Jackson PhD ^104^, Dr JT McCrone PhD ^104^, Dr Nathan Medd PhD ^104^, Dr Emily Scher PhD ^104^ and Jon-Paul Keatley ^116^

**Leadership and supervision:**

Dr Tanya Curran PhD ^3^, Dr Sian Morgan FRCPath ^10^, Prof Patrick Maxwell PhD ^20^, Prof Ken Smith PhD ^20^, Dr Sahar Eldirdiri MBBS, MSc, FRCPath ^21^, Anita Kenyon MSc ^21^, Prof Alison H Holmes MD ^38, 57^, Dr James R Price PhD ^38, 57^, Dr Tim Wyatt PhD ^69^, Dr Alison E Mather PhD ^75^, Dr Timofey Skvortsov PhD ^77^ and Prof John A Hartley PhD ^96^

**Metadata curation:**

Prof Martyn Guest PhD ^11^, Dr Christine Kitchen PhD ^11^, Dr Ian Merrick PhD ^11^, Robert Munn BSc ^11^, Dr Beatrice Bertolusso Degree ^33^, Dr Jessica Lynch MBCHB ^33^, Dr Gabrielle Vernet MBBS ^33^, Stuart Kirk MSc ^34^, Dr Elizabeth Wastnedge MD ^56^, Dr Rachael Stanley PhD ^58^, Giles Idle ^64^, Dr Declan T Bradley PhD ^69, 77^, Nicholas F Killough MSc ^69^, Dr Jennifer Poyner MD ^79^ and Matilde Mori BSc ^110^

**Project administration:**

Owen Jones BSc ^11^, Victoria Wright BSc ^18^, Ellena Brooks MA ^20^, Carol M Churcher BSc ^20^, Mireille Fragakis HND ^20^, Dr Katerina Galai PhD ^20, 70^, Dr Andrew Jermy PhD ^20^, Sarah Judges BA ^20^, Georgina M McManus BSc ^20^, Kim S Smith ^20^, Dr Elaine Westwick PhD ^20^, Dr Stephen W Attwood PhD ^23^, Dr Frances Bolt PhD ^38, 57^, Dr Alisha Davies PhD ^74^, Elen De Lacy MPH ^74^, Fatima Downing ^74^, Sue Edwards ^74^, Lizzie Meadows MA ^75^, Sarah Jeremiah MSc ^97^, Dr Nikki Smith PhD ^109^ and Luke Foulser ^116^

**Samples and logistics:**

Amita Patel BSc ^12^, Dr Louise Berry PhD ^15^, Dr Tim Boswell PhD ^15^, Dr Vicki M Fleming PhD ^15^, Dr Hannah C Howson-Wells PhD ^15^, Dr Amelia Joseph PhD ^15^, Manjinder Khakh ^15^, Dr Michelle M Lister PhD ^15^, Paul W Bird MSc, MRes ^16^, Karlie Fallon ^16^, Thomas Helmer ^16^, Dr Claire L McMurray PhD ^16^, Mina Odedra BSc ^16^, Jessica Shaw BSc ^16^, Dr Julian W Tang PhD ^16^, Nicholas J Willford MSc ^16^, Victoria Blakey BSc ^17^, Dr Veena Raviprakash MD ^17^, Nicola Sheriff BSc ^17^, Lesley-Anne Williams BSc ^17^, Theresa Feltwell MSc ^20^, Dr Luke Bedford PhD ^26^, Dr James S Cargill PhD ^27^, Warwick Hughes MSc ^27^, Dr Jonathan Moore MD ^28^, Susanne Stonehouse BSc ^28^, Laura Atkinson MSc ^29^, Jack CD Lee MSc ^29^, Dr Divya Shah PhD ^29^, Natasha Ohemeng-Kumi MSc ^32, 112^, John Ramble MSc ^32, 112^, Jasveen Sehmi MSc ^32, 112^, Dr Rebecca Williams BMBS ^33^, Wendy Chatterton MSc ^34^, Monika Pusok MSc ^34^, William Everson MSc ^37^, Anibolina Castigador IBMS HCPC ^44^, Emily Macnaughton FRCPath ^44^, Dr Kate El Bouzidi MRCP ^45^, Dr Temi Lampejo FRCPath ^45^, Dr Malur Sudhanva FRCPath ^45^, Cassie Breen BSc ^47^, Dr Graciela Sluga MD, MSc ^48^, Dr Shazaad SY Ahmad MSc ^49, 70^, Dr Ryan P George PhD ^49^, Dr Nicholas W Machin MSc ^49, 70^, Debbie Binns BSc ^50^, Victoria James BSc ^50^, Dr Rachel Blacow MBCHB ^55^, Dr Lindsay Coupland PhD ^58^, Dr Louise Smith PhD ^59^, Dr Edward Barton MD ^60^, Debra Padgett BSc ^60^, Garren Scott BSc ^60^, Dr Aidan Cross MBCHB ^61^, Dr Mariyam Mirfenderesky FRCPath ^61^, Jane Greenaway MSc ^62^, Kevin Cole ^64^, Phillip Clarke ^67^, Nichola Duckworth ^67^, Sarah Walsh ^67^, Kelly Bicknell ^68^, Robert Impey MSc ^68^, Dr Sarah Wyllie PhD ^68^, Richard Hopes ^70^, Dr Chloe Bishop PhD ^72^, Dr Vicki Chalker PhD ^72^, Dr Ian Harrison PhD ^72^, Laura Gifford MSc ^74^, Dr Zoltan Molnar PhD ^77^, Dr Cressida Auckland FRCPath ^79^, Dr Cariad Evans PhD ^85, 109^, Dr Kate Johnson PhD ^85, 109^, Dr David G Partridge FRCP, FRCPath ^85, 109^, Dr Mohammad Raza PhD ^85, 109^, Paul Baker MD ^86^, Prof Stephen Bonner PhD ^86^, Sarah Essex ^86^, Leanne J Murray ^86^, Andrew I Lawton MSc ^87^, Dr Shirelle Burton-Fanning MD ^89^, Dr Brendan AI Payne MD ^89^, Dr Sheila Waugh MD ^89^, Andrea N Gomes MSc ^91^, Maimuna Kimuli MSc ^91^, Darren R Murray MSc ^91^, Paula Ashfield MSc ^92^, Dr Donald Dobie MBCHB ^92^, Dr Fiona Ashford PhD ^93^, Dr Angus Best PhD ^93^, Dr Liam Crawford PhD ^93^, Dr Nicola Cumley PhD ^93^, Dr Megan Mayhew PhD ^93^, Dr Oliver Megram PhD ^93^, Dr Jeremy Mirza PhD ^93^, Dr Emma Moles-Garcia PhD ^93^, Dr Benita Percival PhD ^93^, Megan Driscoll BSc ^96^, Leah Ensell BSc ^96^, Dr Helen L Lowe PhD ^96^, Laurentiu Maftei BSc ^96^, Matteo Mondani MSc ^96^, Nicola J Chaloner BSc ^99^, Benjamin J Cogger BSc ^99^, Lisa J Easton MSc ^99^, Hannah Huckson BSc ^99^, Jonathan Lewis MSc, PgD, FIBMS ^99^, Sarah Lowdon BSc ^99^, Cassandra S Malone MSc ^99^, Florence Munemo BSc ^99^, Manasa Mutingwende MSc ^99^, Roberto Nicodemi BSc ^99^, Olga Podplomyk FD ^99^, Thomas Somassa BSc ^99^, Dr Andrew Beggs PhD ^100^, Dr Alex Richter PhD ^100^, Claire Cormie ^102^, Joana Dias MSc ^102^, Sally Forrest BSc ^102^, Dr Ellen E Higginson PhD ^102^, Mailis Maes MPhil ^102^, Jamie Young BSc ^102^, Dr Rose K Davidson PhD ^103^, Kathryn A Jackson MSc ^107^, Dr Alexander J Keeley MRCP ^109^, Prof Jonathan Ball PhD ^113^, Timothy Byaruhanga MSc ^113^, Dr Joseph G Chappell PhD ^113^, Jayasree Dey MSc ^113^, Jack D Hill MSc ^113^, Emily J Park MSc ^113^, Arezou Fanaie MSc ^114^, Rachel A Hilson MSc ^114^, Geraldine Yaze MSc ^114^ and Stephanie Lo ^116^

**Sequencing and analysis:**

Safiah Afifi BSc ^10^, Robert Beer BSc ^10^, Joshua Maksimovic FD ^10^, Kathryn McCluggage Masters ^10^, Karla Spellman FD ^10^, Catherine Bresner BSc ^11^, William Fuller BSc ^11^, Dr Angela Marchbank BSc ^11^, Trudy Workman HNC ^11^, Dr Ekaterina Shelest PhD ^13, 81^, Dr Johnny Debebe PhD ^18^, Dr Fei Sang PhD ^18^, Dr Sarah Francois PhD ^23^, Bernardo Gutierrez MSc ^23^, Dr Tetyana I Vasylyeva DPhil ^23^, Dr Flavia Flaviani PhD ^31^, Dr Manon Ragonnet-Cronin PhD ^39^, Dr Katherine L Smollett PhD ^42^, Alice Broos BSc ^53^, Daniel Mair BSc ^53^, Jenna Nichols BSc ^53^, Dr Kyriaki Nomikou PhD ^53^, Dr Lily Tong PhD ^53^, Ioulia Tsatsani MSc ^53^, Prof Sarah O’Brien PhD ^54^, Prof Steven Rushton PhD ^54^, Dr Roy Sanderson PhD ^54^, Dr Jon Perkins MBCHB ^55^, Seb Cotton MSc ^56^, Abbie Gallagher BSc ^56^, Dr Elias Allara MD, PhD ^70, 102^, Clare Pearson MSc ^70, 102^, Dr David Bibby PhD ^72^, Dr Gavin Dabrera PhD ^72^, Dr Nicholas Ellaby PhD ^72^, Dr Eileen Gallagher PhD ^72^, Dr Jonathan Hubb PhD ^72^, Dr Angie Lackenby PhD ^72^, Dr David Lee PhD ^72^, Nikos Manesis ^72^, Dr Tamyo Mbisa PhD ^72^, Dr Steven Platt PhD ^72^, Katherine A Twohig ^72^, Dr Mari Morgan PhD ^74^, Alp Aydin MSci ^75^, David J Baker BEng ^75^, Dr Ebenezer Foster-Nyarko PhD ^75^, Dr Sophie J Prosolek PhD ^75^, Steven Rudder ^75^, Chris Baxter BSc ^77^, Sílvia F Carvalho MSc ^77^, Dr Deborah Lavin PhD ^77^, Dr Arun Mariappan PhD ^77^, Dr Clara Radulescu PhD ^77^, Dr Aditi Singh PhD ^77^, Miao Tang MD ^77^, Helen Morcrette BSc ^79^, Nadua Bayzid BSc ^96^, Marius Cotic MSc ^96^, Dr Carlos E Balcazar PhD ^104^, Dr Michael D Gallagher PhD ^104^, Dr Daniel Maloney PhD ^104^, Thomas D Stanton BSc ^104^, Dr Kathleen A Williamson PhD ^104^, Dr Robin Manley PhD ^105^, Michelle L Michelsen BSc ^105^, Dr Christine M Sambles PhD ^105^, Dr David J Studholme PhD ^105^, Joanna Warwick-Dugdale BSc ^105^, Richard Eccles MSc ^107^, Matthew Gemmell MSc ^107^, Dr Richard Gregory PhD ^107^, Dr Margaret Hughes PhD ^107^, Charlotte Nelson MSc ^107^, Dr Lucille Rainbow PhD ^107^, Dr Edith E Vamos PhD ^107^, Hermione J Webster BSc ^107^, Dr Mark Whitehead PhD ^107^, Claudia Wierzbicki BSc ^107^, Dr Adrienn Angyal PhD ^109^, Dr Luke R Green PhD ^109^, Dr Max Whiteley PhD ^109^, Emma Betteridge BSc ^116^, Dr Iraad F Bronner PhD ^116^, Ben W Farr BSc ^116^, Scott Goodwin MSc ^116^, Dr Stefanie V Lensing PhD ^116^, Shane A McCarthy ^116, 102^, Dr Michael A Quail PhD ^116^, Diana Rajan MSc ^116^, Dr Nicholas M Redshaw PhD ^116^, Carol Scott ^116^, Lesley Shirley MSc ^116^ and Scott AJ Thurston BSc ^116^

**Software and analysis tools:**

Dr Will Rowe PhD^43^, Amy Gaskin MSc ^74^, Dr Thanh Le-Viet PhD ^75^, James Bonfield BSc ^116^, Jennifier Liddle ^116^ and Andrew Whitwham BSc ^116^

**1** Barking, Havering and Redbridge University Hospitals NHS Trust, **2** Barts Health NHS Trust, **3** Belfast Health & Social Care Trust, **4** Betsi Cadwaladr University Health Board, **5** Big Data Institute, Nuffield Department of Medicine, University of Oxford, **6** Blackpool Teaching Hospitals NHS Foundation Trust, **7** Bournemouth University, **8** Cambridge Stem Cell Institute, University of Cambridge, **9** Cambridge University Hospitals NHS Foundation Trust, **10** Cardiff and Vale University Health Board, **11** Cardiff University, **12** Centre for Clinical Infection and Diagnostics Research, Department of Infectious Diseases, Guy’s and St Thomas’ NHS Foundation Trust, **13** Centre for Enzyme Innovation, University of Portsmouth, **14** Centre for Genomic Pathogen Surveillance, University of Oxford, **15** Clinical Microbiology Department, Queens Medical Centre, Nottingham University Hospitals NHS Trust, **16** Clinical Microbiology, University Hospitals of Leicester NHS Trust, **17** County Durham and Darlington NHS Foundation Trust, **18** Deep Seq, School of Life Sciences, Queens Medical Centre, University of Nottingham, **19** Department of Infectious Diseases and Microbiology, Cambridge University Hospitals NHS Foundation Trust, **20** Department of Medicine, University of Cambridge, **21** Department of Microbiology, Kettering General Hospital, **22** Department of Microbiology, South West London Pathology, **23** Department of Zoology, University of Oxford, **24** Division of Virology, Department of Pathology, University of Cambridge, **25** East Kent Hospitals University NHS Foundation Trust, **26** East Suffolk and North Essex NHS Foundation Trust, **27** East Sussex Healthcare NHS Trust, **28** Gateshead Health NHS Foundation Trust, **29** Great Ormond Street Hospital for Children NHS Foundation Trust, **30** Great Ormond Street Institute of Child Health (GOS ICH), University College London (UCL), **31** Guy’s and St. Thomas’ Biomedical Research Centre, **32** Guy’s and St. Thomas’ NHS Foundation Trust, **33** Hampshire Hospitals NHS Foundation Trust, **34** Health Services Laboratories, **35** Heartlands Hospital, Birmingham, **36** Hub for Biotechnology in the Built Environment, Northumbria University, **37** Hull University Teaching Hospitals NHS Trust, **38** Imperial College Healthcare NHS Trust, **39** Imperial College London, **40** Infection Care Group, St George’s University Hospitals NHS Foundation Trust, **41** Institute for Infection and Immunity, St George’s University of London, **42** Institute of Biodiversity, Animal Health & Comparative Medicine, **43** Institute of Microbiology and Infection, University of Birmingham, **44** Isle of Wight NHS Trust, **45** King’s College Hospital NHS Foundation Trust, **46** King’s College London, **47** Liverpool Clinical Laboratories, **48** Maidstone and Tunbridge Wells NHS Trust, **49** Manchester University NHS Foundation Trust, **50** Microbiology Department, Buckinghamshire Healthcare NHS Trust, **51** Microbiology, Royal Oldham Hospital, **52** MRC Biostatistics Unit, University of Cambridge, **53** MRC-University of Glasgow Centre for Virus Research, **54** Newcastle University, **55** NHS Greater Glasgow and Clyde, **56** NHS Lothian, **57** NIHR Health Protection Research Unit in HCAI and AMR, Imperial College London, **58** Norfolk and Norwich University Hospitals NHS Foundation Trust, **59** Norfolk County Council, **60** North Cumbria Integrated Care NHS Foundation Trust, **61** North Middlesex University Hospital NHS Trust, **62** North Tees and Hartlepool NHS Foundation Trust, **63** North West London Pathology, **64** Northumbria Healthcare NHS Foundation Trust, **65** Northumbria University, **66** NU-OMICS, Northumbria University, **67** Path Links, Northern Lincolnshire and Goole NHS Foundation Trust, **68** Portsmouth Hospitals University NHS Trust, **69** Public Health Agency, Northern Ireland, **70** Public Health England, **71** Public Health England, Cambridge, **72** Public Health England, Colindale, **73** Public Health Scotland, **74** Public Health Wales, **75** Quadram Institute Bioscience, **76** Queen Elizabeth Hospital, Birmingham, **77** Queen’s University Belfast, **78** Royal Brompton and Harefield Hospitals, **79** Royal Devon and Exeter NHS Foundation Trust, **80** Royal Free London NHS Foundation Trust, **81** School of Biological Sciences, University of Portsmouth, **82** School of Health Sciences, University of Southampton, **83** School of Medicine, University of Southampton, **84** School of Pharmacy & Biomedical Sciences, University of Portsmouth, **85** Sheffield Teaching Hospitals NHS Foundation Trust, **86** South Tees Hospitals NHS Foundation Trust, **87** Southwest Pathology Services, **88** Swansea University, **89** The Newcastle upon Tyne Hospitals NHS Foundation Trust, **90** The Queen Elizabeth Hospital King’s Lynn NHS Foundation Trust, **91** The Royal Marsden NHS Foundation Trust, **92** The Royal Wolverhampton NHS Trust, **93** Turnkey Laboratory, University of Birmingham, **94** University College London Division of Infection and Immunity, **95** University College London Hospital Advanced Pathogen Diagnostics Unit, **96** University College London Hospitals NHS Foundation Trust, **97** University Hospital Southampton NHS Foundation Trust, **98** University Hospitals Dorset NHS Foundation Trust, **99** University Hospitals Sussex NHS Foundation Trust, **100** University of Birmingham, **101** University of Brighton, **102** University of Cambridge, **103** University of East Anglia, **104** University of Edinburgh, **105** University of Exeter, **106** University of Kent, **107** University of Liverpool, **108** University of Oxford, **109** University of Sheffield, **110** University of Southampton, **111** University of St Andrews, **112** Viapath, Guy’s and St Thomas’ NHS Foundation Trust, and King’s College Hospital NHS Foundation Trust, **113** Virology, School of Life Sciences, Queens Medical Centre, University of Nottingham, **114** Watford General Hospital, **115** Wellcome Centre for Human Genetics, Nuffield Department of Medicine, University of Oxford, **116** Wellcome Sanger Institute, **117** West of Scotland Specialist Virology Centre, NHS Greater Glasgow and Clyde, **118** Whittington Health NHS Trust

**Supplementary Table 1.** A review of published studies investigating the impact of viral variant on Ct values.

| Reference | Ct values |  |  |  | How were individuals chosen to be part of the study | Longitudinal? | How was strain determined? |
| --- | --- | --- | --- | --- | --- | --- | --- |
|  | Predecessor variants to B.1.17 | B.1.1.7 | B.1.351 | Delta |  |  |  |
| Frampton [12] | Ct Mean=32<br>SD=4.8<br>N=143 | Mean Ct=28.8<br>SD=4.7<br>N=341 |  |  | Individuals acutely admitted to hospital in London. | No | S-gene target failure |
| Calistri [13] | Median Ct=16.9<br>95% CI=[10.4,19.9]<br>N=965 | Median Ct=15.8<br>95% CI=[9.6,19.6]<br>N=313<br>p-value<10<sup>-4</sup> (relative to predecessor) |  |  | Swabs from three provinces of Abruzzo in Italy were collected based on clinical symptoms or reported contact with confirmed COVID-19 cases. | Yes, but only from some individuals. Infected periods were calculated from only those individuals with 2 or more positive samples. | Whole genome sequencing |
| Kidd [14] | median Ct (ORF1ab) =22.30<br>median Ct (N gene)=23.1<br>N=450 | median Ct (ORF1ab)=18.16<br>median Ct (N gene)=19.39<br>N=178<br>p-value<10<sup>-5</sup> (relative to predecessor for both ORF1ab and N gene) |  |  | Samples in the UK Department of Health and Social Care Test and Trace network. | No | S-gene target failure |
| Cosentino [15] | Ct at self reported symptom onset=22.7<br>95% CI=[22.4,23.0]<br>N=3272 | Ct at self reported symptom onset= 21.3<br>95% CI=[21.1,21.6]<br>N=11496<br>p-value<10<sup>-6</sup> (relative to predecessor) | Ct at self reported symptom onset= 21.6<br>95% CI=[21.1,22.0]<br>N=1366<br>p-value<10<sup>-6</sup> (relative to predecessor) |  | Community testing. Symptomatic individuals | Principally no, although 20% of individuals had two or more samples | S-gene target failure |
| Kissler [16] | Peak viral concentration: Ct=20.1<br>95% CI=[18.3,21.7]<br>N=41 | Peak viral concentration Ct=21.0<br>95% CI=[19.1,20.9]<br>N=36<br>No meaningful difference compared to predecessor reported |  | Peak viral concentration Ct=19.8<br>95% CI=[18.0,22.0]<br>N=36<br>No meaningful difference compared to predecessor reported | Individuals associated with a professional basketball league | Yes (testing done daily) | Whole genome sequencing |
| Ke [17] | Predicted minimum Ct (in saliva)= 23.7<br>N=44 | Predicted minimum Ct (in saliva)= 24.2<br>N=16<br>p-value=0.32 (relative to predecessor) |  |  | Positive samples, or contacts of positive samples, from twice weekly testing of all faculty, staff and students at a university campus. | Yes (daily testing for up to 14 days) | Whole genome sequencing |
| Pouwels [11] |  | Median Ct=31.6 *<br>IQR=[22.8,33.7]<br>N=577 |  | Median Ct=30.1<br>IQR=[18.6,33.7]<br>N=110 | Population representative survey (includes asymptomatic) | Yes, although samples are typically spaced 1 week or 1 | Estimated based upon sample date |
|  |  | *in new PCR positives<br>vaccinated for 14<br>≥21 days after dose 1 or<br><14 days after dose 2 |  | in new PCR-positives<br>vaccinated for 14<br>≥21 days after dose 1 or<br><14 days after dose 2 |  | month apart,<br>meaning that<br>most recorded<br>infections have<br>only one<br>positive sample |  |
| Li [19] | Median Ct =34.31<br>IQR=[31-36]<br>N=63 |  |  | Median Ct =24<br>IQR=[19-29]<br>N=62 | Close contacts of confirmed cases | Yes (daily<br>testing) | Whole genome<br>sequencing |

**Supplementary table 2.** Variance inflation factor (VIF) values.

|  | Variance inflation factor (VIF) |  |
| --- | --- | --- |
|  | Oxford | Northumbria |
| **Sample date** | 7.5* | 3.4* |
| **Age** | 1.6 | 2.8 |
| **Prior exposure**<br>*Ref= vaccine and AB -ve* |  |  |
| AB +ve | 2.1 | 4.1 |
| Vaccinated | 4.1 | 5.2 |
| **Variant (in vaccine and AB -ve)**<br>*Ref=Alpha (Oxford), Delta(Northumbria)* |  |  |
| B.1.177 | 1.1 |  |
| Delta | 1.2 |  |
| BA.1 Omicron |  | 1.3 |
| **Variant (in AB+ve)**<br>*Ref=Alpha (Oxford), Delta(Northumbria)* |  |  |
| B.1.177 | 1.4 |  |
| Delta | 1.7 |  |
| BA.1 Omicron |  | 4.1 |
| **Variant (in vaccinated)**<br>*Ref=Alpha (Oxford), Delta(Northumbria)* |  |  |
| B.1.177 | 1.0 |  |
| Delta | 5.0 |  |
| BA.1 Omicron |  | 2.2 |
| **Vaccine dose in vaccinated**<br>*Ref=1 dose* |  |  |
| 2 doses | 2.3 | 5.3 |
| 3 doses |  | 2.1 |
| **Sex**<br>*Ref=female* |  |  |
| Male | 1.0 | 1.0 |
| **Vaccine product**<br>*Ref=Pfizer* |  |  |
| AstraZeneca | 1.4 | 1.5 |

**Supplementary Figure 1.**
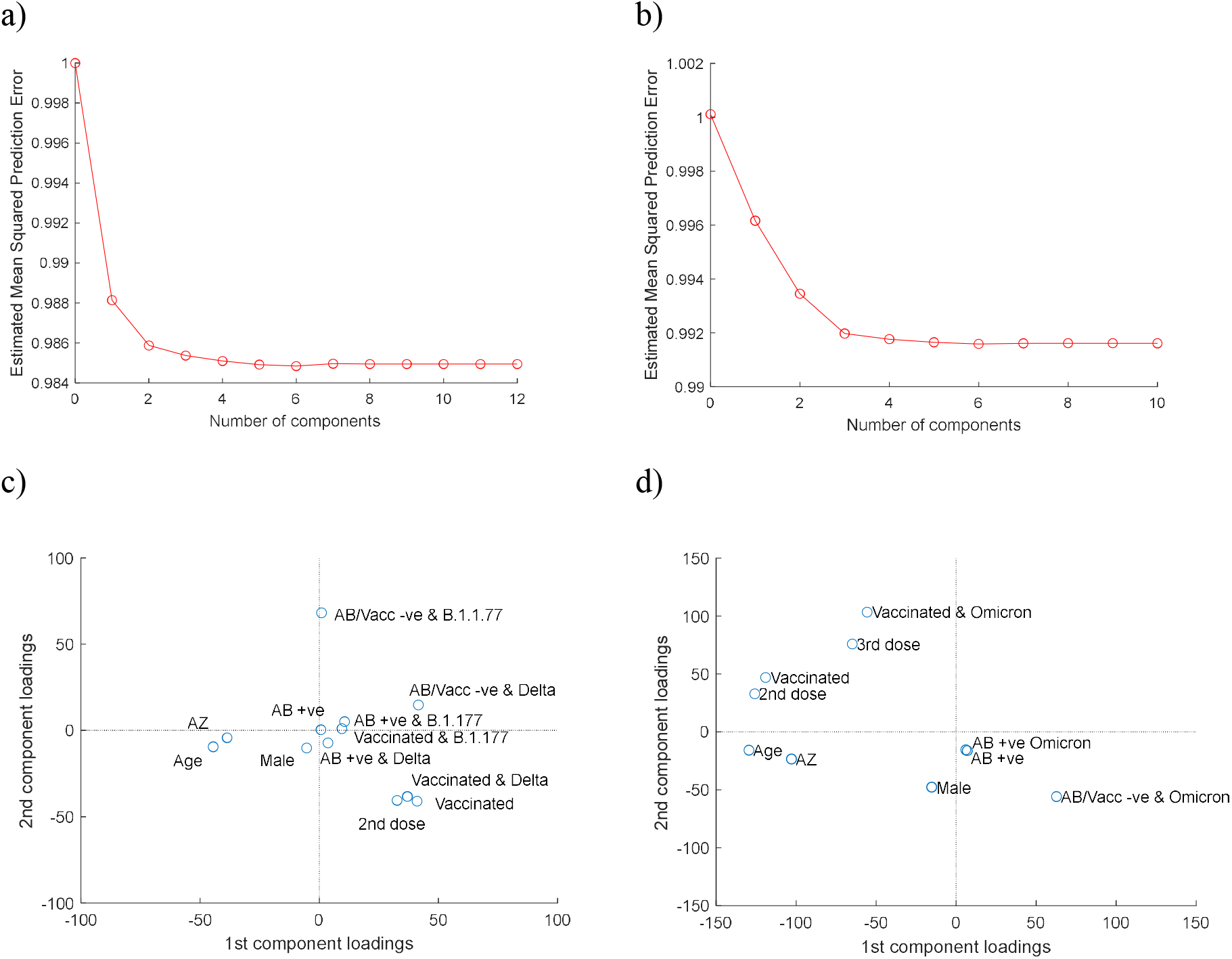
Mean squared error and loading plots relating to the partial least squares regression analysis. a) The mean squared error (MSE) plot for the samples sequenced at Oxford show that 6 latent components minimise the MSE. b) The mean squared error plot for the samples sequenced at Northumbria show that 6 latent components minimise the MSE. c) The loading plot relating to the first two latent components for the samples sequenced at Oxford show that age, vaccine product, vaccination status and variant (Delta vs Alpha) amongst vaccinated and unvaccinated individuals most strongly to the first component; and that variant (B.1.177 vs Alpha) amongst individuals with no known prior exposure contributes most strongly to the second component. d) The loading plot relating to the first two latent components for the samples sequenced at Northumbria show that age, vaccination status and vaccine product contribute most strongly to the first component; and variant (BA.1 Omicron vs Delta) amongst vaccinated individuals contributes most strongly to the second component.

